# An RNA Degradation Complex Required for Spreading and Epigenetic Inheritance of Heterochromatin

**DOI:** 10.1101/870766

**Authors:** Gergana Shipkovenska, Alexander Durango, Marian Kalocsay, Steven P. Gygi, Danesh Moazed

**Affiliations:** Howard Hughes Medical Institute, Department of Cell Biology, Harvard Medical School, Boston, MA, USA; Department of Cell Biology, Harvard Medical School, Boston, MA, USA

## Abstract

Heterochromatin assembly requires the methylation of histone H3 lysine 9 by the Clr4(Suv39h) methyltransferase and both the spreading and epigenetic inheritance of heterochromatin involve the recognition of H3K9me-containing nucleosomes by Clr4 and catalysis of H3K9me on adjacent nucleosomes. How this read-write mechanism overcomes obstacles posed by RNA polymerase II and nascent RNA in its path is not fully understood. Here we identify a role for the highly conserved and essential Rix1-containing complex (here referred to as the rixosome), with known RNA endonuclease and polynucleotide kinase activities required for ribosomal RNA (rRNA) processing, in spreading and epigenetic inheritance of heterochromatin. Viable mutations in rixosome subunits that disrupt its association with Swi6/HP1 fail to localize to heterochromatin, lead to accumulation of heterochromatic RNAs, and block spreading of H3K9me and silencing away from nucleation sites into actively transcribed regions. These findings reveal a new pathway for degradation of heterochromatic RNAs with essential roles in heterochromatin spreading and inheritance.

## Introduction

Heterochromatic domains of DNA are a conserved feature of eukaryotic chromosomes and play central roles in regulation of transcription, inactivation of transposons, and maintenance of genome integrity (Allshire and Madhani, 2018, Saksouk et al., 2015). Heterochromatin is established by the recruitment of histone-modifying enzymes to nucleation sites, followed by spreading of the modification by a read-write mechanism, which involves recognition of the initially deposited modification and deposition of the same modification on histones in adjacent nucleosomes. The so-formed broad domains of repression can be epigenetically inherited through many cell divisions. Recent studies have demonstrated that both the read-write capability and DNA sequences associated with nucleation sites are required for epigenetic inheritance of heterochromatin (Audergon et al., 2015, Wang and Moazed, 2017, Yu et al., 2014, Laprell et al., 2017, Coleman and Struhl, 2017, Ragunathan et al., 2015). However, the mechanisms that mediate faithful epigenetic inheritance are not fully understood.

In the fission yeast *Schizosaccharomyces pombe* heterochromatin is assembled at pericentromeric DNA regions, telomeres, the mating type locus, and the ribosomal DNA repeats (Holoch and Moazed, 2015a, Wang et al., 2016). These regions are associated with histone H3K9 methylation, a conserved marker of heterochromatin, which is catalyzed by the Clr4 (Suv39h) methyltransferase (Rea et al., 2000, Nakayama et al., 2001). Studies using an inducible heterochromatin formation system demonstrated that heterochromatin can be inherited independent of any input from underlying DNA sequence (Ragunathan et al., 2015, Audergon et al., 2015). In these studies, H3K9 methylation is induced by fusion of a Clr4 fragment, containing the methyltransferase domain but lacking the chromodomain (CD) required for recognition of H3K9me, to the bacterial Tetracycline Repressor (TetR-Clr4ΔCD), which binds to *tetO* arrays inserted at a euchromatic locus. The inheritance of the inducible heterochromatic domain depends on the read-write capability of Clr4 and is opposed by the anti-silencing factor *epe1*+, which encodes a Jmjc family demethylase member (Trewick et al., 2007, Bao et al., 2019, Sorida et al., 2019). Mutation or deletion of the chromodomain of Clr4, required for efficient read-write, specifically disrupt epigenetic inheritance without any effect on heterochromatin establishment (Ragunathan et al., 2015, Audergon et al., 2015). In wild-type *epe1*+ cells, in addition to the read-write mechanism, epigenetic inheritance of heterochromatin requires input from specific DNA sequences or a locally generated siRNA amplification loop (Wang and Moazed, 2017, Yu et al., 2018). The development of inducible heterochromatin domains, and the demonstration that they could be epigenetically inherited, provides a unique opportunity to delineate the pathways that are specifically required for heterochromatin inheritance.

To investigate whether other pathways work together with the read-write mechanism to maintain heterochromatin, we used the inducible heterochromatin system to conduct a genetic screen for mutations that abolish heterochromatin inheritance without affecting its establishment. Our screen identified mutations in several pathways, including mutations in three subunits of an RNA processing complex, composed of Rix1, Crb3, Grc3, Mdn1, Las1 and Ipi1 (Castle et al., 2012, Schillewaert et al., 2012, Castle et al., 2013, Gasse et al., 2015, Fromm et al., 2017). This complex, which we propose to name the “rixosome”, is conserved from yeast to human, has well-characterized essential functions in ribosomal RNA (rRNA) processing and has been previously shown to localize to heterochromatin in a Swi6-dependent manner (Kitano et al., 2011, Iglesias et al., 2019), but its role in heterochromatin function remains unknown. We show that the rixosome targets heterochromatic RNAs for degradation through the conserved 5’-3’ exoribonuclease pathway to clear a path for Clr4-mediated read-write.

## Results

### Epigenetic maintenance deficient (*emd*) mutants

Since the requirements for establishment and maintenance of heterochromatin cannot be readily separated at endogenous sites, the identification of pathways that may be specifically required for epigenetic inheritance of heterochromatin has not been possible. To identify such pathways, we employed the inducible system which allows for separation of the establishment and maintenance phases of heterochromatin formation (Ragunathan et al., 2015, Audergon et al., 2015)(Figure 1A). After heterochromatin establishment, the TetR-Clr4ΔCD fusion is released by the addition of tetracycline to disrupts the interaction of the TetR domain with the *10xtetO* targeting sequence. The maintenance of silencing is then monitored by the color of the colonies grown on tetracycline-containing medium. The *ade6*_OFF_ cells give rise to red colonies, due to the accumulation of a red intermediate in the adenine biosynthesis pathway, while the *ade6*_ON_ cells give rise to white colonies. In the absence of the anti-silencing factor Epe1 (*epe1Δ*), growth on tetracycline-containing medium results in a variegated phenotype, due to the stable propagation of the heterochromatic domain in roughly 60% of cells (Audergon et al., 2015, Ragunathan et al., 2015).

**Figure 1.**
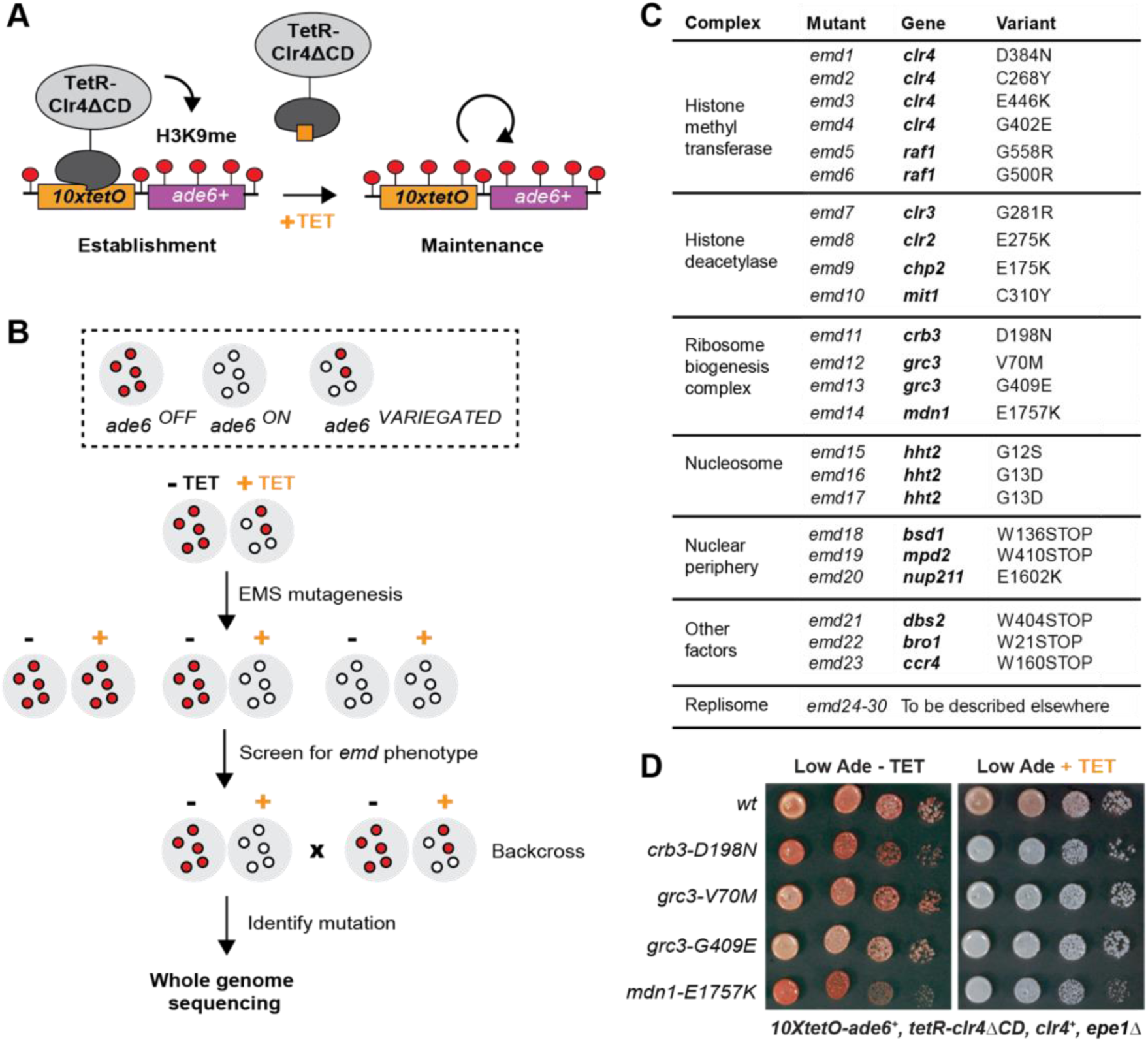
A Genetic Screen Identifies Factors Essential for Heterochromatin Inheritance. (**A**) Schematic diagram representing an inducible reporter system for heterochromatin establishment and inheritance (Ragunathan et al., 2015, Audergon et al., 2015). *10xtetO*, 10 copies of the *tet* operator; TetR-Clr4ΔCD, fusion of TetR and Clr4 lacking the chromodomain; H3K9me, di- and tri-methyl histone H3 Lys 9; TET, tetracycline. (**B**) Schematic diagram of the screen design. EMS, ethyl methanesulfonate; *emd*, epigenetic maintenance deficient. (**C**) List of mutations identified by whole genome sequencing, with their respective genes and the protein complexes that they are known to associate with, where applicable. (**D**) Silencing assay on low adenine medium in the absence (-TET) or presence (+TET) of tetracycline for mutants *emd11-14,* carrying point mutations in subunits of the rixosome. Ade, adenine.

We performed random mutagenesis by ethyl methanesulfonate (EMS) treatment of cells carrying this reporter and screened for *epigenetic maintenance defective* (*emd*) mutants (Figure 1B). We plated the mutagenized cultures on low adenine medium lacking tetracycline and then replica plated the mutant colonies on low adenine medium containing tetracycline (Figure S1A). We screened the resulting replicas for colonies which lost the silencing of the *10xtetO-ade6*+ reporter in the presence of tetracycline, but not in its absence. Such colonies represent mutants that are specifically defective for maintenance of silencing, because they are proficient in heterochromatin establishment mediated by the TetR-Clr4ΔCD initiator, but cannot propagate the silent state after the release of the initiator from the reporter. We screened 80,000 colonies and obtained 83 apparent *emd* mutants (Figure S1B). About half of these mutants showed single-locus segregation patterns in random spore analysis, when backcrossed to the original reporter strain. The remaining mutants showed complex segregation patterns, consistent with cumulative weak effects from multiple contributing loci. We proceeded to characterize the *emd* mutants with single gene segregation patterns, as these likely represent the factors with the strongest effects on heterochromatin maintenance.

While all *emd* mutants lost silencing on tetracycline-containing medium, they differed in their ability to establish efficient silencing on medium lacking tetracycline (Figure S1C). In one class of mutants, the silencing of the *10xtetO-ade6*+ reporter was slightly weaker than that of the non-mutagenized strain, indicating that the mutated genes likely play a role in heterochromatin establishment, in addition to their role in maintenance. In the remaining mutants, the establishment of silencing was either as strong as or stronger than that of the non-mutagenized reporter strain, suggesting that the underlying mutations are specifically required for heterochromatin maintenance. The stronger than wild-type establishment may reflect weakened heterochromatin structure at the endogenous domains and the resulting redistribution of silencing factors from the endogenous to the ectopic domain.

To isolate the causative variants in each of the *emd* mutants, we performed whole genome sequencing combined with pooled linkage analysis (Birkeland et al., 2010, Iida et al., 2014). Out of the 40 *emd* mutants, 30 yielded high confidence single nucleotide mutations (the mutation was present in >90% of the mutant spore pool), one had no mutation despite good genome coverage, and the remaining 9 had insufficient sequencing coverage to be analyzed. The identified mutations mapped to 19 genes, most of which encode subunits of one of five complexes (Figure 1C). Among these are the CLRC and SHREC complexes, which play key roles in heterochromatin formation, and novel mutations in genes with no known function in heterochromatin maintenance. These included three independently identified mutations in the N-terminal tail of histone H3, encoded by *hht2,* and multiple mutations in different components of the DNA replication machinery (to be reported elsewhere). In addition, we isolated mutations in multiple genes, encoding factors broadly associated with the nuclear periphery (*bsd1*, *nup211*, *mpd2*), a subunit of the major Ccr4-Not deadenylase complex (ccr4), a universal stress family protein (*dbs2*), and a vacuolar sorting protein (*bro1*). Finally, we identified four mutations in three genes – *crb3*, *grc3* and *mdn1 –* which are subunits of an essential rRNA processing and ribosome biogenesis complex that we refer to as the rixosome (Figure 1D) and are the focus of this study.

The rixosome contains six unique subunits – three structural subunits Crb3, Rix1 and Ipi1, which form the core of the complex, and three catalytic subunits – the endonuclease Las1, the polynucleotide kinase Grc3 and the AAA-type ATPase Mdn1 (Figure 2A)(Fromm et al., 2017). All subunits are essential for viability and are conserved from yeast to mammals, including humans (Figure 2A). Our screen identified four mutations in three different subunits of the complex – Crb3, Grc3 and Mdn1. We validated three of the mutations by reconstituting them *de novo* in the original reporter strain. All *de novo* constructed strains reproduced the mutant phenotypes in plating assays on low adenine medium in the presence and absence of tetracycline (Figure 2B). To confirm that the observed maintenance defects were due to loss of heterochromatin, we followed the propagation of the heterochromatin specific histone H3 Lys9 di-methylation (H3K9me2) in the absence of tetracycline and in cultures treated with tetracycline for 10 generations. In the absence of tetracycline, the rixosome mutants showed similar levels of H3K9me2 as the wild-type, which is consistent with the lack of silencing defects under the establishment conditions (Figure 2C). By contrast, in the tetracycline treated cultures, the levels of H3K9me2 were drastically reduced in the mutants, but not in the wild-type, indicating that loss of silencing of the *10xtetO-ade6*+ reporter under the maintenance conditions resulted from loss of the underlying heterochromatin (Figure 2D).

**Figure 2.**
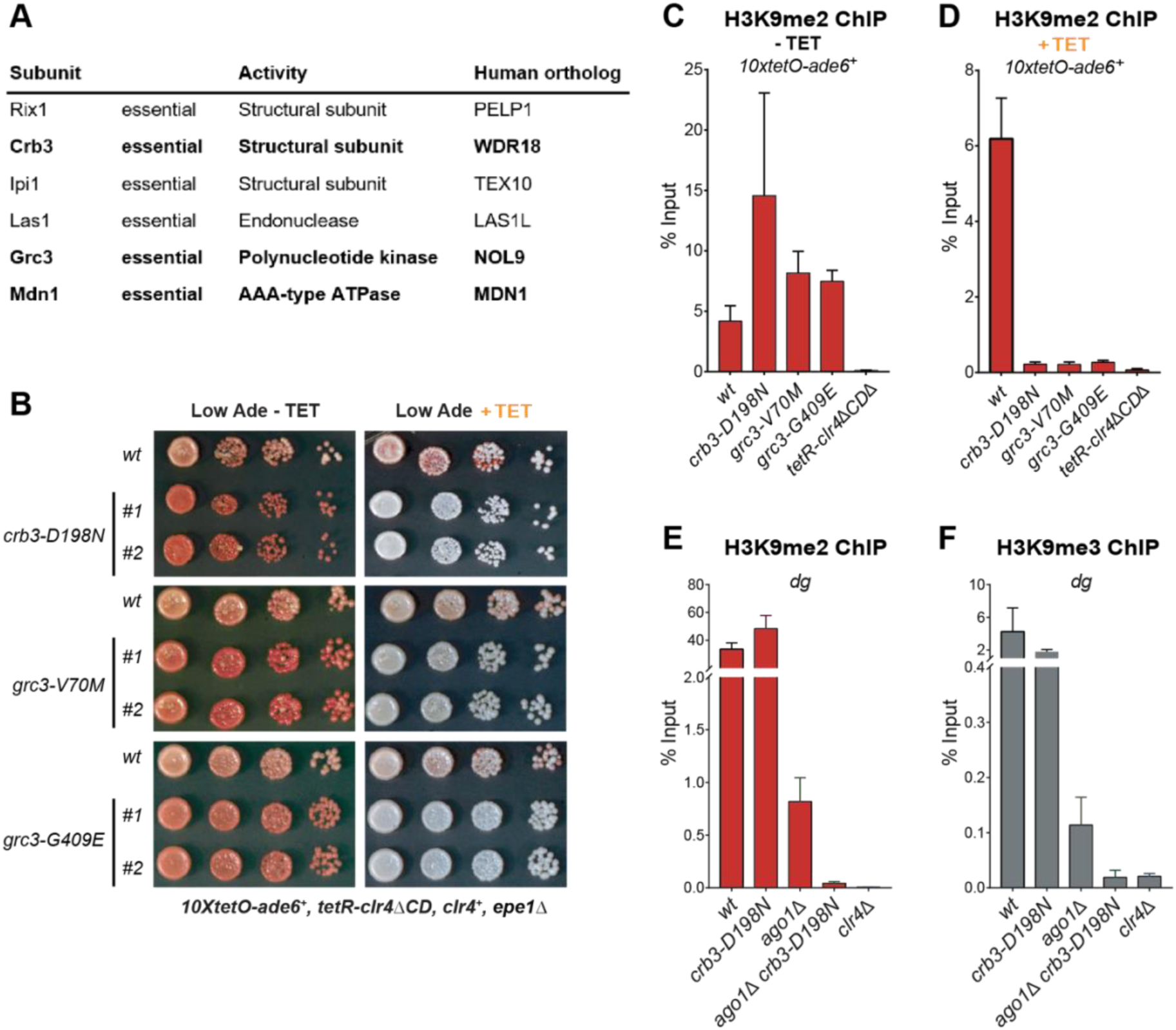
The Rixosome is Required for the Inheritance of Heterochromatin. (**A**) Subunits of the rixosome, their specific activities in the context of ribosome biogenesis, and their human orthologs. Subunits identified in our screen are highlighted in bold. (**B**) Silencing assays to validate the phenotypes of mutants derived from the screen. Three mutations in rixosome subunits were reconstituted *de novo* in *10xtetO-ade6+* reporter cells and plated on low adenine medium in the presence or absence of tetracycline to assess heterochromatin establishment and maintenance. (**C, D**) Chromatin immunoprecipitations (ChIPs) for di-methylated histone H3 Lys 9 (H3K9me2) at the *10xtetO-ade6+* reporter under establishment and maintenance conditions in wild-type cells and cells carrying mutations in rixosome subunits. (**E, F**) ChIP for H3K9me2 and H3K9me3 at the pericentromeric *dg* repeats. Mutations in the rixosome subunit *crb3* abolish H3K9me2 and me3 in the absence of ongoing establishment by the RNAi pathway.

We next investigated whether the rixosome mutants also cause silencing defects at the endogenous heterochromatic domains. To assess if the mutations affect silencing at the pericentromeric and the mating type locus repeats, we introduced the *crb3-D198N* allele in strains carrying the *ura4*+ transgene inserted inside the centromeric *otr1R* and the mating type *cenH* regions, respectively. We then assayed for *ura4*+ silencing by plating cells on minimal medium lacking uracil (-Ura) or rich medium containing 5-fluoroacetic acid (5FOA). Uracil supplementation is essential for cell growth in the absence of *ura4*+ expression, while 5FOA is toxic to cells expressing *ura4*+. Thus, growth on -Ura and 5FOA medium can be used as a readout of *ura4*+ expression. In the presence of ongoing establishment from the siRNA generating *otr1R* and *cenH* repeats, silencing at these regions was robust in mutant cells (Figure S2). Both the mutant and wild-type grew poorly on -Ura medium. At moderate concentrations of 5FOA (0.1%) the growth of the mutant was comparable to that of the wild-type and only at high concentrations (0.2% 5FOA), in which cells are sensitive to very low levels of *ura4*+ expression, weak growth defects could be detected (Figure S2). These results indicate that the establishment of heterochromatin at endogenous domains is largely unaffected by the *crb3-D198N* mutation, consistent with the phenotype at the ectopic reporter.

Is the rixosome required for the maintenance of H3K9me at the endogenous heterochromatic domains? At the pericentromeric DNA regions, siRNAs from the *dg* and *dh* repeats are required for efficient H3K9 methylation and heterochromatin formation (Volpe et al., 2002). Mutations in the RNAi pathway, such as deletion of Ago1, reduce the levels of H3K9me2 and me3, but do not eliminate them (Sadaie et al., 2004). The remaining H3K9 methylation levels reflect ongoing heterochromatin maintenance requiring the Clr4 read-write mechanism (Ragunathan et al., 2015). We introduced the *crb3-D198N* mutation in the *ago1*+ and *ago1Δ* cells and compared the levels of H3K9me2 and me3 at the pericentromeric *dg* repeats to the levels observed in *clr4Δ* cells, which lack histone H3K9 methylation (Figure 2E, F). The *crb3-D198N* mutant did not significantly reduce the levels of H3K9me2 or me3 at the *dg* repeats compared to the *crb3*+ strain, consistent with the results in the *10xtetO-ade6*+ reporter locus (Figure 2B, C, D), which showed no silencing defects in the presence of ongoing establishment. By contrast, in *ago1Δ* cells, where establishment is abolished, H3K9me2 and me3 were reduced to background levels by the *crb3-D198N* mutation (Figure 2E, F). These results show that the rixosome is required for the maintenance of H3K9me at endogenous heterochromatic domains.

Since the rixosome plays an essential role in ribosome biogenesis, we tested whether the rixosome mutations isolated in our screen affect rRNA processing. rRNA precursors accumulate in mutants which disrupt processing and can be detected by gel electrophoresis. In a known rRNA processing mutant in the Crb3 subunit of the rixosome (Kitano et al., 2011), rRNA precursors accumulate at elevated temperatures (Figure S3). We observed weak rRNA processing defects at elevated temperatures in cells carrying *crb3-D198N* and *grc3-G409E* mutations. By contrast, no rRNA precursors accumulated in the *grc3-V70M* mutant (Figure S3), which we therefore conclude is a separation of function allele best suited for further analysis of rixosome function in heterochromatin formation.

### A mutation that abolishes rixosome heterochromatic localization

To determine whether the rixosome mutations affect the integrity of the complex or its interactions with other proteins, we affinity purified wild-type and mutant rixosome complexes and analyzed them by quantitative tandem mass tag mass spectrometry (TMT-MS)(Navarrete-Perea et al., 2018). The rixosome co-purifies with native heterochromatin fragments (Iglesias et al., 2019). We therefore used this native purification protocol to test whether the rixosome mutations affect its interaction with native heterochromatin (Iglesias et al., 2019). We tagged the C-terminus of the Crb3 subunit with a tandem affinity purification (TAP) tag (Puig et al., 2001), which did not interfere with the maintenance of silencing at the *10xtetO-ade6*+ reporter (Figure S4A). We then purified the rixosome using its Crb3-TAP subunit from wild-type, Crb3-D198N, or Grc3-V70M cell extracts (Figure 3A, Figure S4B, C) and analyzed the purifications by TMT-MS. The Crb3-TAP purification from wild-type cells pulled down the core subunits reported previously to be part of the rixosome (Figure 3B, Figure S4D) and its rRNA processing and ribosome biogenesis interaction network (Tables S3 and S4). Importantly, we also recovered the heterochromatin-associated protein Swi6, which was previously shown to associate with the rixosome in an H3K9me-independent manner (Iglesias et al., 2019). When we purified Crb3-TAP from cells carrying the *crb3-D198N* mutation, we detected a modest reduction in the amount of co-precipitated Swi6 in the mutant (Figure S4D), which was not explained by a decrease in the overall Swi6 protein levels (Figure S4E). However, other interactions, including those between the subunits of the rixosome complex (Figure S4D, Table S4), were also weakly affected, indicating that the stability of the complex in the *crb3-D198* mutant may have been compromised. These results were consistent with the mild rRNA processing defects observed in this mutant at elevated temperatures (Figure S3A). By contrast, when we purified Crb3-TAP from cells carrying the *grc3-V70M* mutation, the levels of Swi6 copurifying with the mutant complex were dramatically reduced, but the mutation had little or no effect on the overall integrity of the rixosome and its other interactions (Figure 3B, Table S3). Furthermore, the *grc3-V70M* mutation had no effect on the overall levels of Swi6 in the cell, ruling out indirect effects due to reduced Swi6 expression or stability (Figure 3C). These results demonstrate that the Grc3-V70M mutation specifically disrupts the association of the rixosome complex with Swi6 and further support a specific role for the complex in heterochromatin maintenance, independent of its essential function in ribosome biogenesis. The Grc3-V70M mutation is located in the unstructured N-terminus of Grc3 within a VxVxV sequence motif, a variant of the consensus PxVxL motif of proteins, which interact directly with the chromoshadow domain of HP1 proteins (Figure 3D)(Thiru et al., 2004). The substitution of the central valine in this motif, which is crucial for the interaction between HP1 and HP1-interacting proteins (Thiru et al., 2004), with methionine, would be expected to disrupt the binding surface. Therefore, Swi6 likely interacts directly with the N-terminus of Grc3 via its chromoshadow domain.

**Figure 3.**
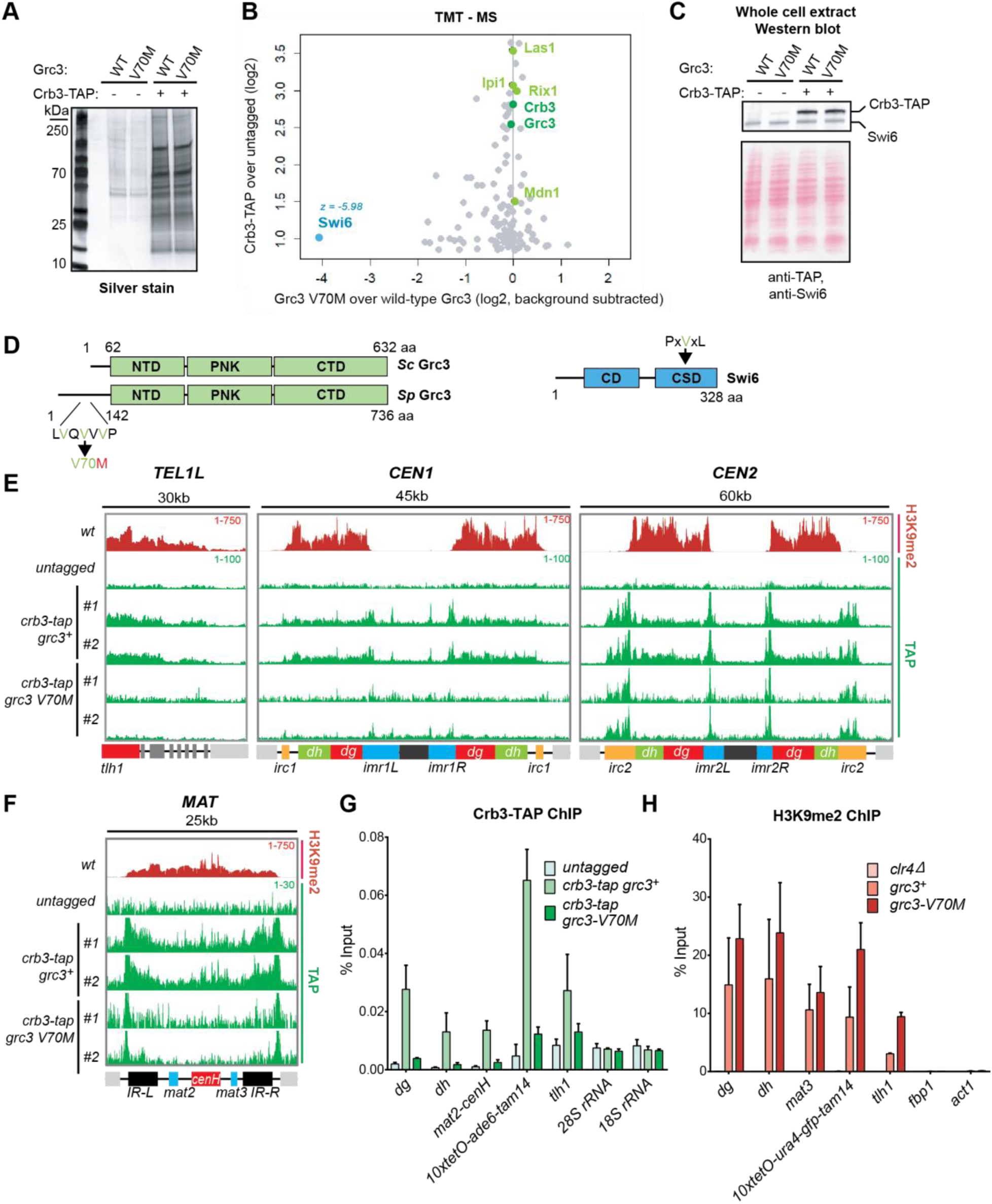
A mutation in the rixosome Grc3 subunit abolishes its interaction with the HP1 protein Swi6 and releases the mutant complex from heterochromatin. (**A**) Silver staining of immunoprecipitations of wild-type and *grc3-V70M* mutant rixosome complexes from cell extracts. (**B**) Tandem mass tag mass spectrometry (TMT-MS) of the immunoprecipitations in A. Subunits of rixosome highlighted in green, HP1 protein Swi6 in blue, ribosome biogenesis and other factors in gray. For detailed description of TMT-MS data analysis see the STAR Methods section. (**C**) Top, western blot from whole cell extracts for the rixosome subunit Crb3-TAP and the HP1 protein Swi6 from *grc3+* and *grc3-V70M* cells. Bottom, ponceau staining is shown as a loading control. (**D**) Schematics of domain organizations of Grc3 (left) and Swi6 (right). Homologous regions of *S.p.* Grc3 and *S. cerevisiae* (S.*c.)* Grc3 are indicated (Pillon et al., 2017). The extended N-terminus of S.*p.* Grc3 lacks similarity to *S.c.* Grc3. The position and amino acid context of the V70M mutation are shown. The three residues highlighted in green are a variant of a consensus motif (PxVxL), recognized by the chromoshadow domain of HP1 proteins. NTD, N-terminal domain; PNK, polynucleotide kinase domain; CTD, C-terminal domain; CD, chromodomain; CSD, chromoshadow domain. (**E, F**) ChIP sequencing profiles of H3K9me2 and Crb3-TAP at heterochromatin in *grc3+* and *grc3-V70M* cells. *tel*, telomere; *cen*, centromere; *mat*, mating type locus; kb, kilobases. Numbers in top right corner of each panel show reads per million. For detailed notations, see Figure S2 legend. (**G, H**) ChIP-qPCR quantification of rixosome and H3K9me2 enrichment at different heterochromatic regions in wild-type and *grc3-V70M* mutant cells. Error bars represent standard deviations for three biological replicates. *dg* and *dh*, tandem repeats in pericentromeric DNA repeats; *tlh1*, subtelomeric gene; *act1* and *fbp1*, euchromatic controls.

We next asked whether the Grc3-V70M and Crb3-D198N mutations affect the localization of the complex to heterochromatic DNA regions. We performed chromatin immunoprecipitations (ChIPs) of Crb3-TAP from wild-type, *crb3-D198N* and *grc3-V70M* cells, followed by sequencing or qPCR. As shown in Figure 3E, F and G, Crb3-TAP co-localized with H3K9me2-containing heterochromatic domains. In the *crb3-D198N* mutant, which only partially reduced the association of the complex with the heterochromatic protein Swi6, the localization of the rixosome was only modestly reduced (Figure S5A). This result suggests that the impairment of epigenetic maintenance in *crb3-D198N* mutant cells results from a rixosome defect downstream of its localization to heterochromatic domains. By contrast, in the *grc3-V70M* mutant, heterochromatin localization of the Crb3-TAP was reduced to background levels (Figure 3E, F and G, Figure S5B). Furthermore, Crb3-TAP localized to the ectopically induced domain of H3K9 methylation at the *tetO-ade6*+ reporter and this localization was abolished in *grc3-V70M* cells (Figure S5C). Finally, Crb3-TAP was not enriched at the ribosomal DNA locus or the subtelomeric *tel1R and tel2L* regions in wild-type cells, despite the presence of H3K9me2 (Figure S5D). In addition to heterochromatic regions, the rixosome showed enrichment at a number of euchromatic loci, which did not contain H3K9me2 and were not affected by the *grc3-V70M* mutant (Figure S5E), including tRNA genes. Consistent with unperturbed heterochromatin establishment in *grc3-V70M* mutant cells (Figure 1D, Figure 2B), the mutation did not affect the levels of H3K9me2 at various heterochromatic loci (Figure 3H). These results suggest that the failure of rixosome complexes, carrying the Grc3-V70M mutant subunit, to localize to heterochromatin results in defective epigenetic maintenance.

### The rixosome promotes Dhp1/XRN2-mediated RNA degradation

In the context of rRNA processing in *S. cerevisiae*, the endonuclease and polynucleotide kinase activities of the rixosome cleave the rRNA precursor and prepare the resulting RNA ends for downstream processing (Castle et al., 2012, Castle et al., 2013, Schillewaert et al., 2012, Gasse et al., 2015). The processing of the 3’ end is carried out by the exosome, in a molecular hand-off event, where the exosome subunit Dis3 removes the 2’-3’ cyclic phosphate produced by the Las1 endonucleolytic cleavage and hands off the 3’end to the Rrp6 subunit for further degradation (Fromm et al., 2017). The processing of the 5’ end is carried out by the nuclear 5’-3’ exonuclease Dhp1, a homolog of the budding yeast Rat1 and human XRN2 proteins, which recognizes the free 5’ phosphate group deposited by Grc3 (Gasse et al., 2015). Moreover, in *S. pombe*, Dhp1 has previously been shown to promote transcription termination within heterochromatin and a *dhp1* mutation impairs the silencing of some heterochromatic reporter genes, although how heterochromatic RNAs become substrates for Dhp1 had remained unknown (Chalamcharla et al., 2015, Tucker et al., 2016). We hypothesized that the rixosome is specifically recruited to heterochromatin to initiate the degradation of heterochromatic RNAs, by channeling them through the 5’-3’ and 3’-5’ exonucleases Dhp1 and Dis3/Rrp6, respectively. To test this hypothesis, we introduced the previously described *dhp1-1* (Shobuike et al., 2001), and *dis3-54* (Murakami et al., 2007, Ohkura et al., 1988) mutant alleles into cells carrying the *10xtetO-ade6*+ reporter. The *dhp1-1* allele had no effect on silencing of the *10xtetO-ade6*+ gene in the absence of tetracycline, under establishment conditions, but abolished silencing in the presence of tetracycline, under maintenance conditions, closely phenocopying the rixosome mutants (Figure 4A). Consistent with the silencing results, chromatin immunoprecipitation showed that H3K9me2 was lost in *dhp1-1* cells, in the presence of tetracycline but not in its absence (Figure 4B). Relative to the *dhp1-1* allele, the *dis3-54* allele disrupted maintenance to a lesser extent (Figure S6A). Together, these results suggest that 5’-3’ mediated RNA degradation by Dhp1, rather than 3’-5’ mediated RNA degradation by the exosome, is the key event downstream of the rixosome required for epigenetic inheritance.

**Figure 4.**
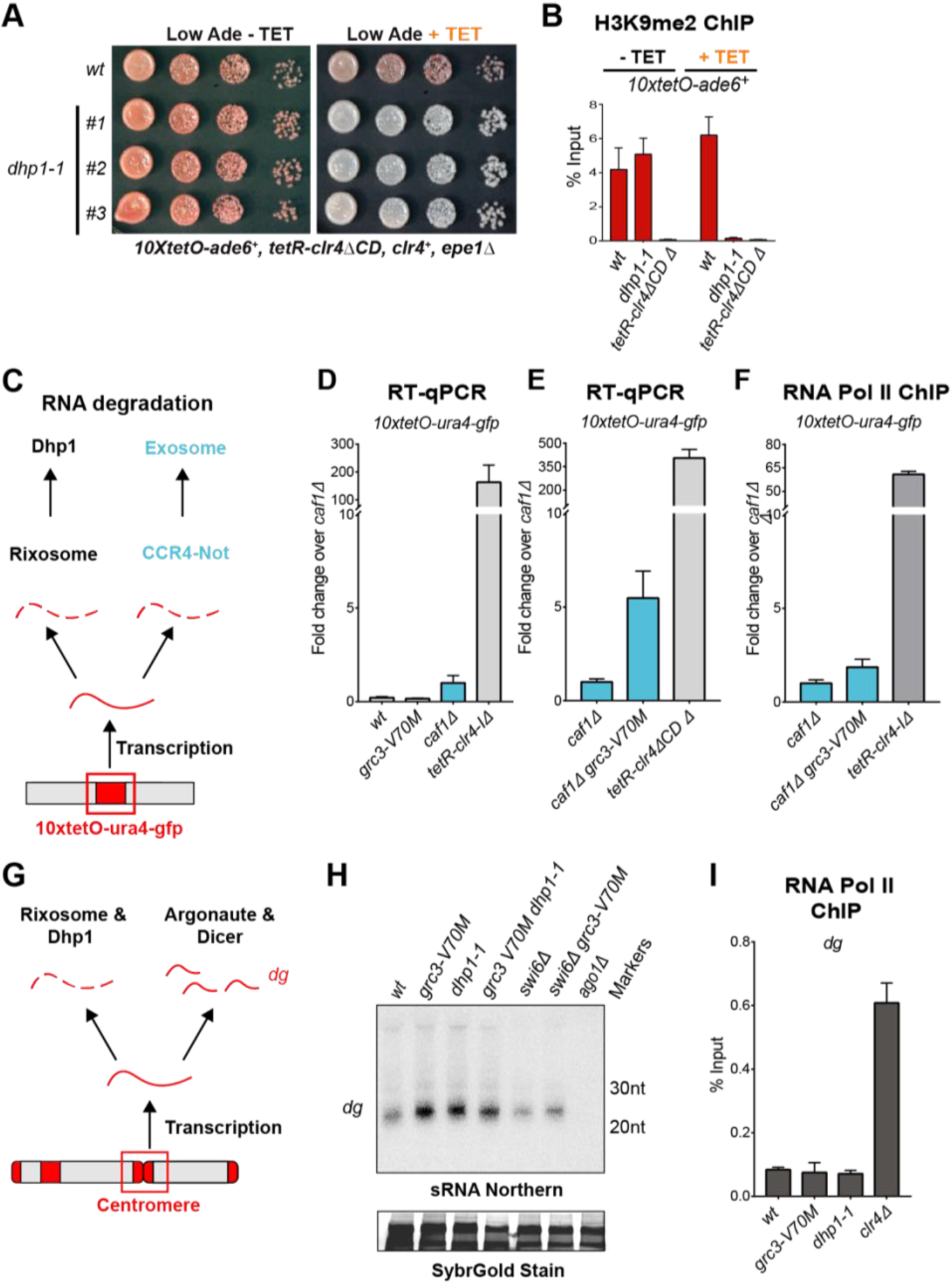
The rixosome channels heterochromatic RNAs for degradation by the 5’-3’ exonuclease Dhp1/Xrn2. (**A**) Silencing assay for heterochromatin establishment and maintenance at the *10xtetO-ade6+* reporter locus in a Dhp1/Xrn2 hypomorphic mutant, *dhp1-1*. (**B**) ChIP for H3K9me2 at the *10xtetO-ade6+* reporter locus under establishment and maintenance conditions in cells carrying *dhp1+* or *dhp1-1* alleles. (**C**) Model for the role of the rixosome in RNA degradation at the *10xtetO-ura4-gfp/ade6+* reporter. The rixosome cleaves RNAs produced from the reporter on heterochromatin and targets them for degradation by the nuclear Xrn2/Dhp1 exonuclease. If an RNA escapes degradation by the rixosome, it is deadenylated by the Ccr4-Not complex, in which Caf1 is a key subunit, in preparation for decapping and degradation by the cytoplasmic exosome. Deletion of the deadenylation pathway unmasks the effect of the rixosome and Xrn2/Dhp1 on reporter RNAs. (**D, E**) Reverse transcription (RT) -qPCR assay for RNA accumulation at the *10xtetO-ura4-gfp* reporter locus in wild-type, *grc3-V70M* and *caf1Δ* single mutants (**D**) and in *caf1Δ and caf1Δ grc3* double mutant (**E**). Caf1 is a subunit of the major cytoplasmic RNA deadenylase complex, Ccr4-Not. (**F**) ChIP for RNA polymerase II (Pol II) at the reporter locus in *caf1Δ and caf1Δ grc3-V70M*. (**G**) Model for the role of the rixosome in RNA degradation at the pericentromeric repeats. The rixosome acts on full length RNAs transcribed from the repeats in competition with RNAi and targets them for degradation through the exonuclease Xrn2/Dhp1. When the rixosome is released from heterochromatin or when Dhp1 activity is reduced, *dg* transcripts normally degraded by the rixosome and Dhp1 are channeled for processing by RNAi, resulting in increased siRNA accumulation. (**H**) Small RNA Northern blot, measuring the accumulation of small RNAs originating from the pericentromeric *dg* repeats in wild-type and different mutants. (**I**) ChIP for RNA polymerase II (Pol II) at the pericentromeric *dg* repeats.

To investigate whether the effects of the rixosome and the exonuclease Dhp1 on heterochromatin are indeed mediated via RNA degradation, we measured the levels of heterochromatic RNAs in the *grc3-V70M* and *dhp1-1* mutant cells. To test the effects of the mutations on the steady-state levels of RNAs transcribed from the *10xtetO* reporter locus, we used the previously described *10xtetO-ura4*+-*gfp* version of the reporter (Ragunathan et al., 2015). Unlike *ade6*, which is present in the genome in two copies, an ectopic wild-type copy and endogenous copy carrying a point mutation, the *gfp* transgene presents a unique sequence for quantitative reverse transcription (qRT) and RNA Pol II ChIP. In initial experiments, under establishment condition, we did not observe an increase in *gfp* reporter RNA levels in the *grc3-V70M* mutant relative to wild-type cells. We reasoned that other redundant RNA degradation pathways may mask the effects of the rixosome on RNA levels under strong establishment conditions at the reporter locus. To test this idea, we deleted a component of the nuclear TRAMP complex (*cid14Δ*) and a component of the Ccr4-Not complex (*caf1Δ*)(LaCava et al., 2005, Buhler et al., 2007, Tucker et al., 2001), the other major RNA degradation pathways previously shown to impact heterochromatin (Figure 4C). TRAMP channels nuclear RNAs for degradation to the exosome, while the Ccr4-Not complex is the principal deadenylase, which prepares transcripts for decapping and degradation in the cytoplasm by the exosome and Xrn1, (Tucker et al., 2001, LaCava et al., 2005). Deletion of *cid14*+ did not affect heterochromatin maintenance and the *cid14Δ grc3-V70M* double mutant cells did not show any increase in *gfp* RNA relative to *cid14Δ* alone (Figure S6B, C). By contrast, the *caf1Δ grc3-V70M* double mutant showed a marked accumulation of *gfp* transcripts relative to *caf1Δ* alone (Figure 4D and E). The increase in RNA levels in the double mutant was not accompanied by a corresponding increase in RNA Pol II occupancy (Figure 4F). Overall, we conclude that heterochromatic RNAs derived from the *10xtetO-ura4*+-*gfp* reporter locus are targeted for degradation by the rixosome and by the Ccr4-Not complex on chromatin and the cytoplasm, respectively (Figure 4C).

We next examined whether rixosome impairment affects the levels of RNAs transcribed from the pericentromeric DNA repeats. RNAs transcribed from these repeats are processed by the RNAi pathway into siRNAs. We reasoned that since the rixosome localizes to these pericentromeric repeats through association with Swi6, it might act in parallel with RNAi to degrade RNAs transcribed from the repeats. If this were the case, we would expect an increase in the level of siRNAs in the absence of the rixosome as more RNA would be available for processing by the RNAi machinery (Figure 4G). We assessed the levels of siRNA generated from the pericentromeric *dg* repeats in wild-type, *grc3-V70M,* and *dhp1-1* cells by northern blotting and, consistent with the above hypothesis, found that both mutations caused a marked increase in siRNAs accumulation (Figure 4H, lanes 1-3). Moreover, the *grc3-V70M dhp1-1* double mutant had siRNA levels comparable to those of each of the single mutants, suggesting that the rixosome and the exonuclease Dhp1 act in the same pathway (Figure 4H, lane 4). Furthermore, *dg* siRNA levels were similar between *swi6Δ grc3*+ and *swi6Δ grc3-V70M* cells, indicating that the ability of the rixosome pathway to compete with RNAi required its Swi6-mediated recruitment (Figure 4H, lanes 5 and 6). Finally, the increased accumulation of pericentromeric dg siRNAs in the above mutant cells was not accompanied by increased RNA polymerase II (RNA Pol II) occupancy at the *dg* repeats (Figure 4I), suggesting that the mutations did not affect the levels of transcription from the repeats. We therefore propose that the rixosome cleaves and phosphorylates heterochromatic RNAs so that they become substrates for co-transcriptional degradation by the Dhp1 exonuclease. Furthermore, this pathway appears to act in parallel with the RNAi pathway at the pericentromeric repeats.

### Role for the rixosome in heterochromatin spreading

To investigate whether mutations in the rixosome or Dhp1 affect the stability of heterochromatic domains genome-wide, we performed ChIP for H3K9me2 and me3, followed by sequencing or qPCR, in wild-type and mutant cells. While the pattern or levels of H3K9me2 were generally not perturbed in *grc3-V70M* and *dhp1-1* mutants relative to wild-type cells, the mutant cells displayed a modest reduction in H3K9me3 levels. This reduction only occurred at heterochromatic domains where the rixosome was enriched: the *mat* locus (Figure S7A and B), the pericentromeric repeats of *cen1*, *cen2* and *cen3* (Figure S7A-D) and the *10xtetO-ade6*+ reporter under establishment conditions (Figure S7A). Since the read-write function of Clr4 is required for the transition from H3K9me2 to me3, these results raise the possibility that the rixosome-Dhp1 pathway is required for efficient Clr4 read-write.

In addition to efficient H3K9me3 deposition and epigenetic inheritance, Clr4 read-write is required for RNAi-independent spreading of H3K9me away from nucleation sites. We therefore tested whether the RNAi-independent spreading of heterochromatin at the *L(BglII):ade6*+ reporter (Ayoub et al., 1999) at the mating type locus was rixosome- and Dhp1-dependent (Figure 5A). In wild-type cells, heterochromatin spreads from the RNAi initiating centromere Homology (*cenH*) region into the *mat2p* gene and the adjacent *L* region. Transgenes inserted into the *L* region display variable levels of silencing due to stochastic spreading of H3K9 methylation and can therefore be used as a readout for heterochromatin spreading (Figure 5B, top row). As a control, we first tested the effects of the read-write deficient *clr4-W31G* mutant on *L(BglII):ade6*+ silencing. As expected, the *L(BglII):ade6*+ reporter became fully derepressed in *clr4-W31G* cells, indicating that Clr4 read-write was essential for the spread of silencing away from the *cenH* nucleation region (Figure 5B, bottom two rows). We then replaced the wild-type *grc3*+ and *dhp1*+ genes with the *grc3-V70M* and *dhp1-1* mutant alleles in cells carrying the *L(BglII):ade6*+ reporter. Both mutants abolished the silencing of the *ade6*+ reporter, in a manner similar to the *clr4-W31G* mutant (Figure 5C, D). Furthermore, analysis of RNA levels using qRT-PCR showed strong accumulation of *ade6*+ RNA in the *grc3-V70M* and *dhp1-1* mutant cells relative to wild-type cells, to an equal or greater extent as that observed in *clr4Δ* cells (Figure 5E). Surprisingly, ChIP-qPCR experiments showed that neither *grc3-V70M* and *dhp1-1* mutations nor *clr4* mutations (*clr4-W31G* and *clr4Δ*) caused an increase in RNA Pol II occupancy (Figure 5F), despite maximal derepression of *cenH* and *ade6*+ RNAs (Figure 5E). Consistent with previous reports (Zhang et al., 2008), the levels of H3K9me2 and H3K9me3 were strongly reduced in *clr4-W31G* cells and did not spread outside the *cenH* region (Figure 5G and H). In *grc3-V70M* and *dhp1-1* cells, H3K9me2 spread efficiently from the *cenH* region into the silent *mat2* and *mat3* cassettes, but failed to spread over the actively transcribed *ade6*+ reporter gene (Figure 5G and H). Importantly, the *grc3-V70M* and *dhp1-1* mutants showed efficient spreading of heterochromatin into the *L* region in the absence of the reporter locus, as evident from the H3K9me2 and me3 ChIP-seq profiles (Figure S7B).These results uncover an essential role for the rixosome in spreading of H3K9me over and silencing of a transcription unit that is inserted distal to a heterochromatin nucleation center, and further indicate that this silencing occurs at a step following transcription.

**Figure 5.**
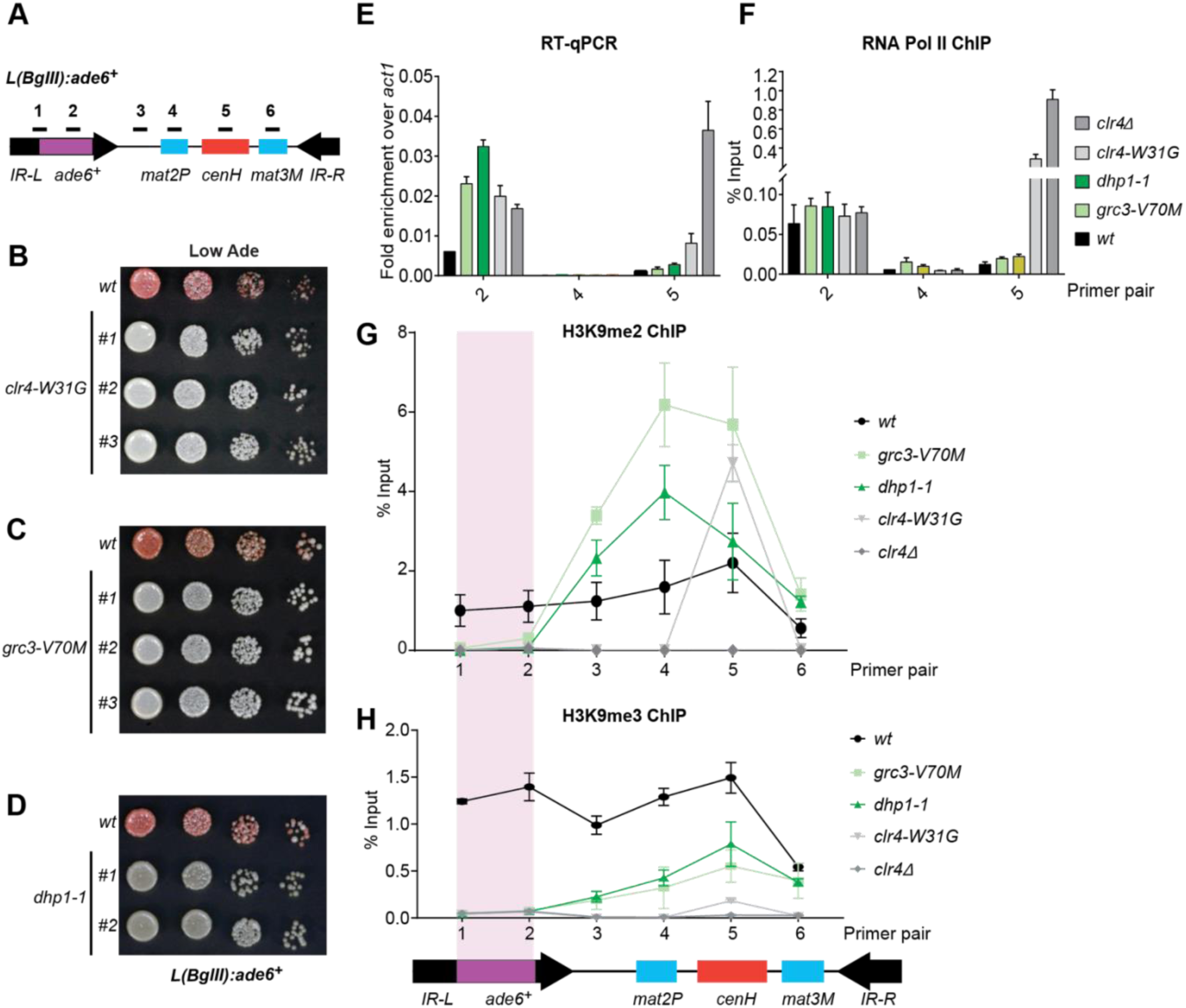
Rixosome and Dhp1 are required for RNAi-independent heterochromatin spreading and promote the Clr4-catalyzed transition H3K9me2 to H3K9me3. (**A**) Schematic of DNA sequence organization of the mating type locus with position of the *L(BglII):ade6+* reporter and centromere homology (*cenH*, red) region highlighted. For detailed notations, see Figure S2 legend. (**B, C, D**) Silencing assays of the *L(BglII):ade6+* reporter in wild type, *clr4-W31G, grc3-V70M* and *dhp1-1* cells. (**E**) Qunatitative Reverse Transcription PCR (qRT) for transcripts along the mating type locus in wild type cells and cells carrying *clr4-W31G*, *grc3-V70M*, *dhp1-1*, and *clr4Δ* mutant alleles. (**F**) ChIP-qPCR for RNA Polymerase II occupancy along the mating type locus in wild type cells and cells carrying *clr4-W31G*, *grc3-V70M*, *dhp1-1*, and *clr4Δ* mutant alleles. (**G, H**) ChIP-qPCR for H3K9me2 and H3K9me3 at the *L(BglII):ade6+* reporter and surrounding regions at the mating type locus in wild type cells and cells carrying *clr4-W31G*, *grc3-V70M*, *dhp1-1*, and *clr4Δ* mutant alleles.

## Discussion

Our ability to perform a sensitive genetic screen for mutations that specifically abolish heterochromatin maintenance allowed us to isolate separation of function alleles and demonstrate a vital role for the essential ribosome biogenesis complex, the rixosome, in epigenetic inheritance of heterochromatin and spreading of H3K9me along the chromatin fiber. The rixosome is recruited to heterochromatin via an interaction with the HP1 protein, Swi6, to promote the degradation of RNAs transcribed by RNA pol II within heterochromatic domains. In the context of heterochromatin, the rixosome performs a function that is analogous to that of the cleavage polyadenylation specificity factor (CPSF) complex at euchromatic mRNAs (Shi and Manley, 2015). However, unlike CPSF, which is recruited by specific polyadenylation signals in the 3’ ends of mRNAs, the rixosome is recruited throughout heterochromatic domains via association with Swi6/HP1 and targets heterochromatic RNAs for degradation rather than polyadenylation and export. This conclusion is supported by our demonstration that heterochromatic RNAs accumulate in cells carrying rixosome mutations and that a mutation in the 5’-3’ exoribonuclease Dhp1 has the same heterochromatin formation defects as rixosome mutant. Dhp1 is known to act downstream of the rixosome during rRNA processing in the nucleolus and in euchromatin following CPSF-mediated cleavage to promote RNA pol II transcription termination. We propose that in the absence of the rixosome or the RNAi pathway, which also degrades heterochromatic RNAs, nascent transcripts and RNA polymerase II form an obstacle that impedes Clr4 read-write during H3K9me spreading and following DNA replication, when read-write copies H3K9me from parental onto newly deposited histone H3. In support of this hypothesis, localization of the rixosome to heterochromatin is critical for read-write-dependent silencing and spreading of H3K9me into a transcribed transgene insert within heterochromatin. All subunits of the rixosome are conserved from yeast to human, raising the possibility that it functions in chromatin silencing in other eukaryotes.

Our findings suggest a new role for RNA degradation, promoted by recruitment of the rixosome, in formation and epigenetic maintenance of heterochromatin. The rixosome contains subunits with multiple catalytic activities (Castle et al., 2012, Schillewaert et al., 2012, Castle et al., 2013, Gasse et al., 2015, Fromm et al., 2017). These activities, which take place on the pre-60S ribosomal particles, include the Las1-mediated endonucleolytic cleavage of the internal transcribed spacer 2 (ITS2) in 27S pe-rRNA, followed by phosphorylation of the 5’-OH cleavage product to generate a 5’-PO_4_ end, which becomes a substrate for trimming by the 5’-3’ exonuclease Dhp1/XRN2 leading to formation of mature 25S rRNA (Figure S8)(Fromm et al., 2017). The cleaved 3’ end, containing 2’,3’ cyclic phosphate, is trimmed by the 3’-5’ RNA exosome to generate mature 5.8S RNA. Finally, the dynein-related AAA ATPase subunits appears to mechanically remove biogenesis factors from the pre-60S particle to generate 60S ribosomal subunits (Ulbrich et al., 2009). Our findings suggest that, in addition to these essential functions, the rixosome is recruited to heterochromatin where its RNA processing activities are utilized to degrade heterochromatic RNAs in order to promote the spreading and epigenetic inheritance of H3K9me (Figure 6A).

**Figure 6.**
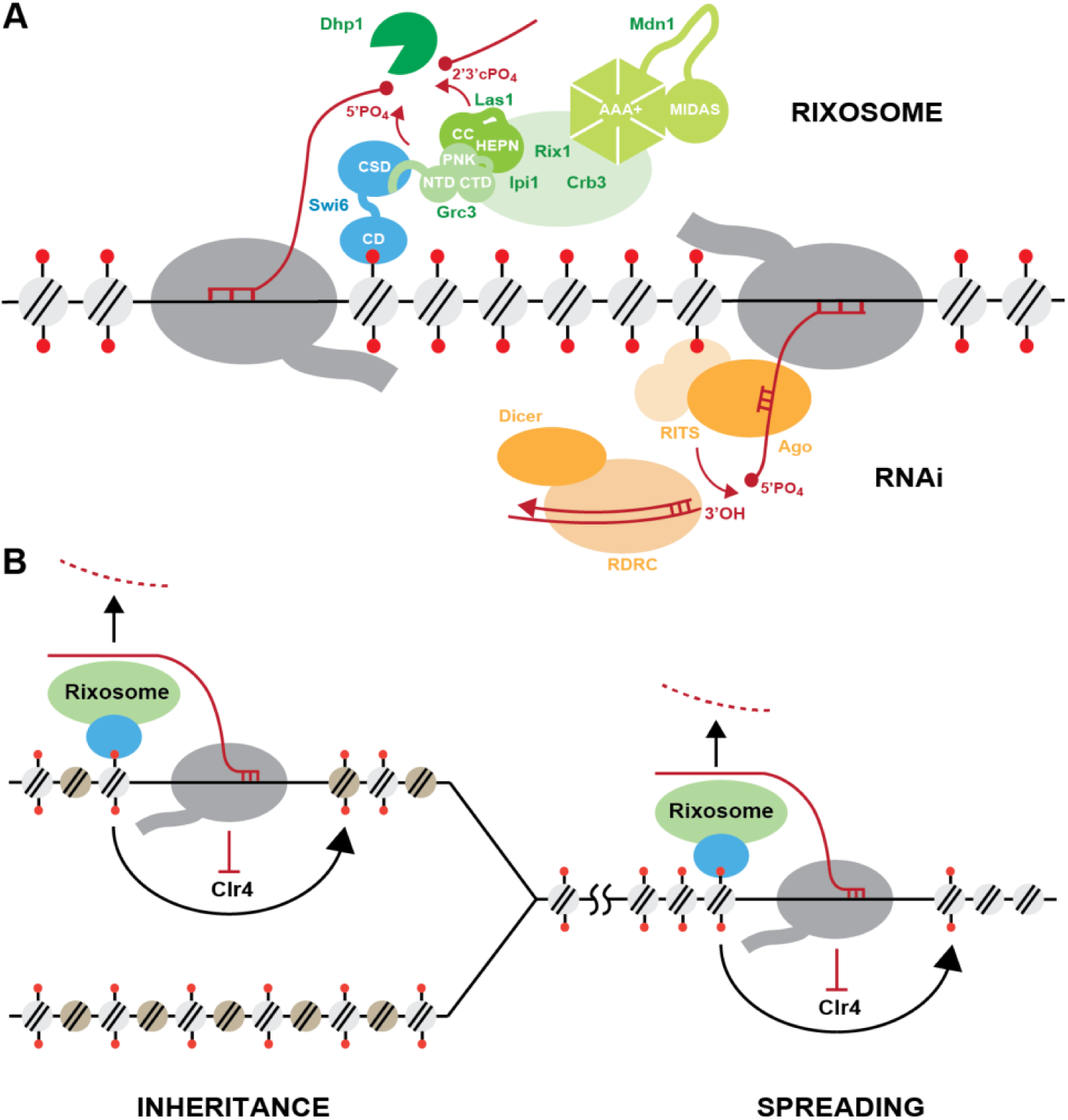
Model for rixosome-mediated degradation of nascent transcripts and its role in spreading and inheritance of heterochromatin. (**A**) Like RNAi, the rixosome localizes to heterochromatin but via a direct interaction between the N-terminus of Grc3 with Swi6, rather than siRNA with the nascent transcript. The endonuclease Las1 and the polynucleotide kinase (PNK) Grc3, subunits of the rixosome, cleave nascent heterochromatic transcripts and phosphorylate the free 5’ hydroxyl end at the site of cleavage, respectively. These processing steps prepare the transcripts for subsequent degradation by the Dhp1/XRN2 exonuclease. CC, coiled coil domain; HEPN, higher eukaryotes and prokaryotes nucleotide-binding domain; AAA+, ATPase domain; Pol II, RNA polymerase II; RITS, RNA-induced Transcriptional Silencing; RDRC, RNA-Dependent RNA Polymerase Complex. (**B**) Nascent RNA clearance enables the spreading and inheritance of heterochromatin by read-write. Nascent transcripts and their associated transcriptional complexes present obstacles in the path of the read-write pathway. By degrading those nascent transcripts, the rixosome and RNAi promote read-write-dependent spreading and inheritance of heterochromatin. In repetitive regions of the genome, where both pathways are present, the RNA clearance functions of RNAi and the rixosome are redundant. In the absence of RNAi, nascent RNA degradation by the rixosome is essential for heterochromatin inheritance and spreading.

The rixosome mutations described in this study abolish the DNA sequence-independent epigenetic inheritance of heterochromatin and, moreover, are completely defective in the spreading of H3K9me from a nucleation region into an inserted transgene. These phenotypes are accompanied by accumulation of heterochromatic RNAs in rixosome mutants. Notably, at pericentromeric DNA repeats where the RNAi machinery is recruited to nascent noncoding RNAs and processes them into siRNAs, the rixosome contributes to noncoding RNA degradation, but is not required for H3K9me spreading or inheritance, unless the RNAi pathway is rendered inactive by mutation. The rixosome thus defines a second RNA degradation mechanism that localizes to heterochromatin, through direct recruitment by Swi6, and plays an indispensable role for heterochromatin spreading and maintenance in transcribed chromosomal regions that are not associated with RNAi (Figure 6B). Both the spreading and epigenetic inheritance of heterochromatin require the read-write capability of the Clr4 methyltransferase. During read-write, Clr4 binds to nucleosomes containing pre-existing H3K9me and catalyzes the methylation of histone H3 in adjacent nucleosomes (Zhang et al., 2008, Al-Sady et al., 2013). The rixosome is likely to cleave nascent pol II-associated RNAs in the path of nucleosome-bound Clr4. In its absence, the nascent RNA pol II complex would create a barrier that blocks Clr4 read-write (Figure 6B), which is required for both spreading from a nucleation point and following DNA replication.

The profound requirement for the rixosome in the spreading and epigenetic inheritance of H3K9me mirrors the requirement for the Clr4 read-write capability and supports the idea that the rixosome and read-write acts coordinately during spreading and inheritance of H3K9me. Furthermore, the rixosome appears to be the key effector that mediates silencing of a transgene inserted at the fission yeast mating type locus. The *ade6*+ transgene inserted at the *L* region of the mating type locus is strongly silenced, but displays similar levels of RNA Pol II occupancy in wild-type and *clr4Δ* cells (Figure 5E, F), indicating that H3K9me-dependent silencing occurs downstream of transcription. The absolute dependence of this silencing on the rixosome and Dhp1 suggests that in this instance silencing occurs co-transcriptionally via RNA degradation. We propose that when genes cannot be silenced at the level of transcription, perhaps due to strong promoters or activation sequences, heterochromatin-localized rixosome triggers the degradation of their transcripts and silences their expression.

In summary, our findings suggest that the rixosome acts as a key trigger that generates substrates for degradation by Dhp1/XRN2 and the above pathways. Finally, we note that subunits of the mammalian rixosome have been reported to interact with chromatin components and mutations in multiple subunits of the complex are associated with human disease (Fanis et al., 2012, Hussey et al., 2014, Sareddy and Vadlamudi, 2016). It will be important to understand the extent to which chromatin-dependent functions of the rixosome may contribute to the regulation of gene expression and development in mammals.

## Materials and Methods

### Yeast strains and growth medium

The genotypes of all strains used in this study are presented in Table S1. Point mutations in the rixosome were reconstituted by seamless integration to avoid confounding effects from the presence of a selection cassette around the target genes. The genes of interest carrying the desired point mutations were cloned into a pFA6a plasmid backbone upstream of a *ura4*+- h*phMX6* cassette, amplified together with the selectable marker and transformed into yeast cells to replace the endogenous gene. The *ura4*+-*hphMX6* cassette was then knocked out by a second transformation to restore the native downstream region. All TAP (tandem affinity purification) tagged strains were expressed under the control of their endogenous promoters and terminators. Cells were cultured on rich YES (yeast extract with supplements) medium, unless otherwise indicated.

### Genetic screen for epigenetic inheritance mutants

Yeast cells carrying the *10xtetO-ade6*+ reporter (Ragunathan et al., 2015) were grown in YES medium with shaking at 32°C to exponential phase (OD600 ∼ 1), treated with 3% ethyl methanesulfonate (EMS)(Winston, 2008) and plated on low adenine YE medium at limiting dilution to form single colonies. The single colonies were replica plated on low adenine medium in the absence (YE) and presence of 5µg/ml tetracycline (YE+TET). The replicas were incubated for 2 days at 32°C and scored for silencing phenotypes. Colonies which displayed silencing on YE medium (red) but not on YE+TET medium (white) were selected for further analysis. After three rounds of single colony purification, the mutants were backcrossed to the original reporter strain and the progeny was subjected to pooled linkage analysis, as described previously (Birkeland et al., 2010). Briefly, crosses were subjected to random spore analysis and 20-24 spores with the mutant phenotype were picked from YE plates. The mutant spores were cultured in individual wells in 24 well plates to saturation, pooled and collected. Genomic DNA was purified with the ZymoResearch Fungal/Bacterial DNA MiniPrep kit and sonicated using QSonica water bath sonicator (15min with 10s ON/OFF pulses at 60% amplitude) to obtain 200-500bp fragment size distribution. Genomic DNA libraries were prepared according to a previously reported protocol (Wilkening et al., 2013) from the original reporter strain and the pools of mutant spores from each cross. Sequencing was performed on an Illumina HiSeq 2500 platform at a depth of at least 3 million reads per mutant pool. Read alignment, variant calling and annotation were performed using the Mudi platform (Iida et al., 2014).

### Growth and silencing assays

Yeast cells were grown in YES medium with shaking at 32°C to stationary phase (OD600 = 12 and above). The cultures were adjusted to a density of OD600 = 4 and serially diluted tenfold. Three microliters of each dilution were spotted on the appropriate medium. For assessing *ura4+* reporter gene silencing assays, serial dilutions were plated on EMMC-Ura (Edinburgh minimal medium lacking uracil) or YES medium supplemented with 5-fluoroorotic acid (0.1% and 0.2% 5FOA). For *ade6+* reporter gene silencing assays, serial dilutions were plated on YE and YE+TET media. Phenotypes were scored after 3 days at 32°C.

### Chromatin immunoprecipitation (ChIP)

For TET treatment, yeast cells were grown for 24h in YES medium supplemented with TET to exponential phase (OD600=1-2). For all other ChIP experiments, yeast cultures were grown in rich (YES) medium to OD600 densities of 1-2. For Crb3-TAP ChIP, 50 OD600 units of cells per ChIP were fixed with 1.5% EGS for 45min and 1% formaldehyde for 15min, quenched with glycine and collected. For all other ChIPs, 50 OD600 units of cells were crosslinked with 1% formaldehyde for 15min. Cell pellets were resuspended in lysis buffer (50mM Hepes/KOH pH 7.5, 140mM NaCl, 1mM EDTA, 1% Triton-X-100, 0.1% Na Deoxycholate, 0.1% SDS, 1mM PMSF, supplemented with protease inhibitors) and cells were lysed by the glass bead method, using MagNA Lyser (Roche) (3 pulses of 90s at 4500rpm, 1 pulse of 45s at 5000rpm). The lysates were sonicated by 3 pulses of 20s (amplitude 40%) in a Branson microtip sonifier with intermittent cooling on ice, to a fragment distribution of 200-1000 bp. Cell debris were removed by centrifugation (16000xg) at 4*C for 15min and the cleared lysates were incubated for 2 hours at 4*C with magnetic beads coupled to antibodies against the targets of interest. For H3K9me2 and RNA Pol II ChIPs, 30µl Protein A Dynabeads (Invitrogen) were coupled with 2µg anti-H3K9me2 antibody (Ab1220, Abcam) or 4µg anti-Pol II (8WG16, Biolegend MMS-126R-500), respectively. For H3K9me3 ChIPs, 30µl Streptavidin Dynabeads M280 (Invitrogen) were coupled to 1µg of anti-H3K9me3 antibody (Diagenode C15500003) and blocked with 5µM biotin. For Crb3-TAP ChIPs, 4.5µg Dynabeads M-270 Epoxy beads (Invitrogen) were coupled to 1.5µg rabbit IgG (Sigma, #15006) and incubated with the cleared lysate. The bead-protein complexes were washed three times with lysis buffer, once with chilled TE buffer (10mM Tris/HCl, 1 mM EDTA) and eluted using 120µl 50mM Tris/HCl pH 8.0, 1mM EDTA, 1% SDS with heating (65oC) for 20min. Crosslinks were reversed at 65°C overnight and the eluates were treated with RNase A and Proteinase K (Roche), prior to an extraction with phenol:chloroform:isoamyl alcohol. qPCR was performed on Applied Biosystems qPCR instrument with the primer sequences listed in Table S2. qPCR quantification and statistical analysis were performed for three biological replicates.

### Library preparation and next generation sequencing

For ChIP-seq, reverse crosslinked DNA treated with RNase A and proteinase K was purified using the Qiagen PCR purification kit. The DNA was then sonicated in a QSonica water bath sonicator (5 min with 15s ON/OFF pulses, 20% amplitude) to a fragment size of ∼200bp, concentrated and quantified using Qubit dsDNA high sensitivity kit. 1-10ng of DNA were used to prepare libraries as described previously (Wong et al., 2013). Libraries were pooled and sequenced on Illumina HiSeq and NextSeq platforms. The raw reads were demultiplexed using Geneious and aligned to the reference genome using bowtie with random assignment of reads to repeats. The mapped reads were normalized to counts per million using Deeptools and visualized in the IGV genome browser.

### Immunoprecipitation of rixosome complexes

Purifications of Crb3-TAP tagged rixosome complexes were performed by a chromatin purification method described previously (Iglesias et al., 2019) with minor modifications. Cells carrying Crb3-TAP in the wild-type and mutant (*grc3-V70M or crb3-D198N*) backgrounds, as well as corresponding untagged cells, were grown in rich YES medium to exponential phase (OD600 = 1-1.5) and 600 OD600 units were collected per immunoprecipitation. The pellets were resuspended in lysis buffer (20 mM Hepes pH7.5-NaOH, 100 mM NaCl, 5 mM MgCl2, 1 mM EDTA pH 8.0, 10% Glycerol, 0.25% Triton-X-100, 0.5mM DTT, 2mM phenylmethylsulphonylfluoride [PMSF], protease inhibitor cocktail). The cells were lysed by the glass bead method in a MagNA Lyser (Roche) with a succession of short pulses (10 cycles, 22s at 5000rpm, 9 cycles, 15s, 5500rpm), followed by rapid cooling for 2min in an ice water bath. Lysates were clarified at 16,000xg for 15min and incubated with 300µl of Pan Mouse IgG Dynabeads (Invitrogen) per IP. Four washes were performed with lysis buffer with full resuspension of the beads, followed by four short washes with wash buffer (20 mM Hepes pH7.5-NaOH, 100 mM NaCl, 5 mM MgCl2). Proteins were eluted with 0.5M NH4OH and dried down in a speed vac. Five to ten percent of each elution fraction was analyzed by silver staining using the Silver Stain SNAP Kit (Pierce) according to the manufacturer’s protocol. The remainder of each elution was analyzed by mass spectrometry.

### Mass spectrometry sample preparation and data analysis

All liquid reagents used for mass spectrometry sample preparation were HPLC grade. Proteins were digested with trypsin (Promega #V5111) in 200mM HEPES buffer pH 8.5 with 2% acetonitrile (v/v) overnight at 37°C. For direct TMT peptide labeling of digests, TMT-10plex reagents from Thermo Fisher Scientific (#90406) were used. After quenching with hydroxylamine at 0.3% (v/v), reactions were acidified with formic acid and peptides purified by reversed phase C_18_ chromatography via stage tips. Data were collected on an Orbitrap Lumos instrument with a multi-notch MS_3_ method. Prior to injection, peptides were separated by HPLC with an EASY-nLC 1000 liquid chromatography system (Thermo Scientific) with a 100μm inner diameter column and a 2.6 μm Accucore C_18_ matrix (Thermo Fisher Scientific) with a 4-hour acidic acetonitrile gradient. MS_1_ scans were recorded in the Orbitrap (resolution 120,000, mass range 400-1400 Th) and MS_2_ scans in the iontrap after collision-induced dissociation (CID, CE=35) and a maximum injection time of 150ms. For TMT quantification, an SPS-MS3 method was used with HDC fragments scanned in the Orbitrap at a resolution of 50,000 at 200 Th with a maximum injection time of 200 ms. Peptides were searched with an in-house software based on SEQUEST (v.28, rev. 12) against a forward and reverse database of the *S. pombe* proteome database (Uniprot) with human contaminants added. Searches had a mass tolerance of 50 ppm for precursors and a MS_2_ fragment ion tolerance of 0.9 Da. For searches, 2 missed cleavages were allowed and methionine oxidation was searched for dynamically (+15.9949 Da). After linear discriminant analysis (LDA), peptide false discovery rate was 1%, as was the FDR for final collapsed proteins. MS_1_ data were post-search, calibrated, and the search performed again. For TMT quantification, peptides with an isolation specificity of >70% and a summed TMT signal-to-noise (s/n) >200 were filtered. Proteins with at least two peptide quantification events were considered for analysis. For all 1139 quantified proteins, an enrichment score was calculated as the summed TMT signal-to-noise (TMT sum s/n) ratio of the tagged wild-type sample over the untagged TMT sum s/n of the wild-type control. The top 10% most enriched proteins (118) contained all known components of the Rix1 core complex (Crb3/Ipi3, Rix1/Ipi2, Ipi1, Las1, Grc3, Mdn1), many of the pre-60S ribosome particle components (ribosomal and ribosome biogenesis proteins), Swi6 (required for the recruitment of the Rix1 complex to heterochromatin) and nuclear export factors known to be involved in the shuttling of the pre-60S complex to the cytosol. The wild-type/mutant fold change score for each protein was calculated as the ratio of the summed TMT s/n in the wild-type over the mutant IPs, after subtraction of the background from the untagged control and correction for the abundance of the bait protein. The log transformed fold-change scores were plotted against the log transformed enrichment scores to identify proteins which were over- or under-represented in the mutant sample relative to the wild-type. Z scores were calculated from the distribution of fold-change scores of the top 10% most enriched proteins and the Z-scores of outliers were displayed on the plot.

### Western blot

To prepare whole cell protein extracts, yeast cells were grown in YES medium at 32°C with shaking to exponential phase (OD600 = 1) and 25ml of culture (25 OD600 units) were collected. Cells were resuspended in lysis buffer (50mM Tris pH 7.5, 150mM NaCl, 5mM EDTA, 10% glycerol, 1mM PMSF, protease inhibitor cocktail) and lysed by the glass bead method in a MagNA Lyser (Roche) with a succession of short pulses (5 pulses, 45s at 4500 rpm), followed by rapid cooling in an ice water bath. Aliquots of 0.25 and 0.5 OD600 units were resolved by SDS-PAGE electrophoresis, immunoblotted with anti-TAP antibody (PAP, Sigma P1291) and anti-Swi6 antibody (rabbit polyclonal, (Holoch and Moazed, 2015b)), followed by IRDye 700DX-conjugated secondary antibody (LI-COR Biosciences), and imaged on an Odyssey imaging system (LI-COR Biosciences).

### Total RNA extraction and reverse transcription

Yeast cultures were grown to exponential phase (OD600 = 1) and 50 OD600 units of cells were collected. Cells were lysed by the hot phenol method, RNA was purified on-column using RNeasy Mini Kit (Qiagen), treated with Turbo DNase (ThermoFischer) and cleaned up by a second RNeasy Mini kit purification. 500ng of pure RNA were reverse transcribed (Superscript III kit, Invitrogen) using the primers indicated in Table S2 and quantified by qPCR on Applied Biosystems qPCR instrument. *act1+* was used as an internal control. Statistical analysis was performed on three biological replicates.

### rRNA processing

Yeast cultures were grown to exponential phase (OD600 = 1) and 50 OD600 units of cells were collected. Cells were lysed by the hot phenol method, RNA was precipitated with ethanol and resuspended in water. RNA was separated by denaturing gel electrophoresis in a MOPS-formaldehyde buffer system and visualized by staining with EtBr.

### sRNA Northern blot

Yeast cultures were grown in YES medium to exponential phase (OD600 = 1), a cell count was determined by a BrightLine_TM_ hemacytometer and 5×10_7_ cells were collected. sRNAs were purified with the mirVana miRNA isolation kit (Ambion) and 25µg of sRNA were used for Northern blotting as described previously (Buhler et al., 2006). Sequences of oligo probes are listed in Table S2.

## Acknowledgements

We thank Tessi Iida for help and advice in analysis of whole genome sequencing data for identification of mutants, members of the Moazed lab for helpful discussions, Swapnil Parhad, Antonis Tatarakis, Xiaoyi Wang, Andy Yuan, Chen Zhou, and Haining Zhou for comments on the manuscript, and Nahid Iglesias for Swi6 purification protocols. This work was supported by an HHMI summer student internship award (A.D.), a fellowship from the Boehringer Ingelheim Fonds (G.S.), and NIH RO1 GM072805 (D.M.). D.M. is a Howard Hughes Medical Institute Investigator.

