## Supplemental Data for "An RNA Degradation Complex Required for Spreading and Epigenetic Inheritance of Heterochromatin"

#### Supplemental Figure Legends

**Figure S1. Screen for Isolation of Inheritance Defective Mutants.**

(**A**) Representative images of plate replicas on low adenine medium lacking or containing tetracycline. Colonies were tracked between the replicas and those red on -TET and white on +TET plates were selected for further analysis (white arrows).

(**B**) Summary of screen data.

(**C**) Silencing assays for mutants isolated from the screen, together with the top hits recovered by whole genome sequencing. In some mutants, multiple linked variants were detected with high confidence (present in >90% of pooled mutant spores) and are listed together here.

**Figure S2. Heterochromatin Establishment at Endogenous Heterochromatic Domains is Largely Unaffected in Rixosome Mutations.**

(**A**) Top: Schematic diagram of DNA sequence organization of the mating type locus with position of the *Kint2::ura4*$^{+}$ reporter. *cenH*, centromere homology region; *mat2P* and *mat3M*, silent mating type loci; *IR-L* and *IR-R*, boundary elements. Bottom: Silencing assays for *Kint2::ura4*$^{+}$ expression on nonselective medium (N/S), medium lacking uracil (-Ura), or medium supplemented with 0.1% and 0.2% 5-Fluoroorotic acid (5FOA). The higher 5FOA concentration allows the detection of minor defects in heterochromatic silencing.

(**B**) Top: Schematic diagram of DNA sequence organization of the pericentromeric repeats on chromosome 1 and position of the *otr1R:ura4*$^{+}$ reporter. *otr*, outermost repeats; *dg* and *dh*, outermost *dg* and *dh* repeats; *imr*, innermost repeats; *irc*, inverted repeat centromere sequences. Bottom: Silencing assays for *otr1R:ura4*$^{+}$ expression on nonselective medium (N/S), medium lacking uracil (-Ura), or medium supplemented with 5FOA.

**Figure S3. Effects of the rixosome mutants on rRNA processing**

A denaturing gel of total RNA, EtBr staining. Mature rRNA species are labelled as 25S and 18S, rRNA precursors as 27S and 35S. The presence of the higher molecular weight precursors indicates rRNA processing defects.

**Figure S4. Identification of proteins associated with wild-type and *crb3-D198N* mutant complexes.**

(**A**) Silencing assay in cells carrying TAP-tagged or untagged Crb3, showing that the presence of the tag does not disrupt establishment or maintenance of silencing at the *10xtetO-ade6*$^{+}$ reporter locus.

(**B**) Western blot of different fractions from immunoprecipitations of the rixosome. The complex is efficiently solubilized and immunodepleted from native cell extracts, as seen from the amounts of Crb3-TAP in the total, soluble, and unbound fractions.

(**C**) Silver staining of immunoprecipitations of wild-type and *crb3-D198N* mutant rixosome complexes from cell extracts.

(**D**) TMT-MS of the immunoprecipitations in D. Rixosome subunits highlighted in green, HP1 protein Swi6 in blue and other factors in gray.

(**E**) Top, western blot from whole cell extracts with anti-TAP and anti-HP1/Swi6 antibody on *crb3-tap* and *crb3-D198N-tap* whole cell extracts. Bottom, ponceau staining is shown as a loading control.

**Figure S5. Recruitment of rixosome complexes to heterochromatin in wild-type, *crb3-D198N* and *grc3-V70M* cells.**

(**A**) Quantification of rixosome enrichment at different heterochromatic regions in wild-type and *crb3-D198N* mutant cells by ChIP-qPCR. Error bars represent standard deviations of the mean of three biological replicates.

(**B, C, D**) ChIP sequencing profiles of H3K9me2 and Crb3-TAP at heterochromatic loci in *grc3*+ and *grc3-V70M* cells. Numbers in top right corner of each panel denote reads per million. For detailed notations, see Figure S2 legend.

(**E**) ChIP-seq profiles of H3K9me2 and Crb3-TAP at euchromatic regions in *grc3*+ and *grc3-V70M* cells. Numbers in top right corner of each panel denote reads per million.

**Figure S6. TRAMP and the exosome do not affect heterochromatin maintenance.**

(**A**) Silencing assay for heterochromatin maintenance at the *10xtetO-ade6*$^+$ reporter locus in the exosome mutant *dis3-54*.

(**B**) Silencing assay for heterochromatin maintenance at the *10xtetO-ade6*$^+$ reporter locus in cells lacking the Cid14 subunit of the TRAMP complex (*cid14Δ*). The TRAMP complex targets RNAs for degradation by the exosome in the nucleus.

(**C**) RT-qPCR assay for RNA accumulation at the *10xtetO-ura4-gfp* reporter locus for wild-type and *grc3-V70M* mutant rixosome in wild-type and *cid14Δ* cells.  Loss of Cid14 had no effect on the levels of RNAs originating from the *10xtetO-ura4-gfp* reporter.

**Figure S7. The rixosome and Dhp1 are required for efficient H3K9 tri-methylation.**

(**A**) Quantification by ChIP-qPCR of tri-methylation levels at different heterochromatic domains in *grc3-V70M* and wild-type cells. Error bars represent standard deviation for three biological replicates.

(**B, C, D**) ChIP-seq profiles of H3K9me2 and H3K9me3 at the pericentromeric regions of chromosomes 1, 2 and 3 (*cen1*, *cen2* and *cen3*) and the *mat* locus. Numbers in top right corner of each panel denote reads per million. For detailed notations, see Figure S2 legend.

(**E, F**) ChIP-seq profiles of H3K9me2 and H3K9me3 at the subtelomeric loci *tlh1* and *tlh2* in wild-type, *grc3-V70M* and *dhp1-1* cells. Numbers in top right corner of each panel denote reads per million. Panels for H3K9me2 in wild-type cells in (**B**), (**C**), (**E**) and (**F**) use data shown in Figures 3 (**E**) and (**F**), and are shown here for ready comparison.

**Figure S8. Summary of previously described rixosome functions during ribosome biogenesis (Gasse et al., 2015) .**

The 27S rRNA precursor is cleaved by the endonuclease Las1, creating a free 5’ hydroxyl (5’OH) group and a 2’3’ cyclic phosphate (2’3’$PO_4$). The 3’ end is processed by the exosome. The 5’ end is phosphorylated by the polynucleotide kinase activity of Grc3 to produce 5’$PO_4$ and processed by the 5’-3’ exonuclease Dhp1 (XRN2). The AAA+ ATPase Mdn1 remodels the RNA-protein complex to release chaperones to produce mature 60S ribosomal subunits.

**Figure S1. Shipkovenska et al.**

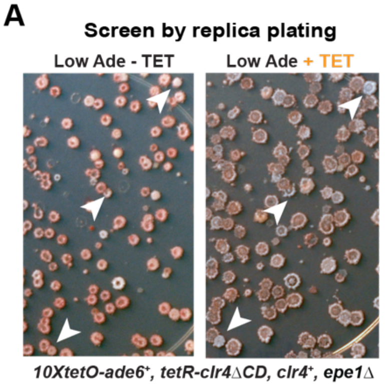


**B** **Summary of screen statistics**

| Screening step | Numbers |
|---|---|
| Total colonies screened | 80,000 |
| Apparent *emd* mutants | 83 |
| *emd* mutants after backcrossing (single gene segregation pattern) | 40 |
| Sequenced (> 3 million reads) | 31 |
| Unique gene hits | 19 |

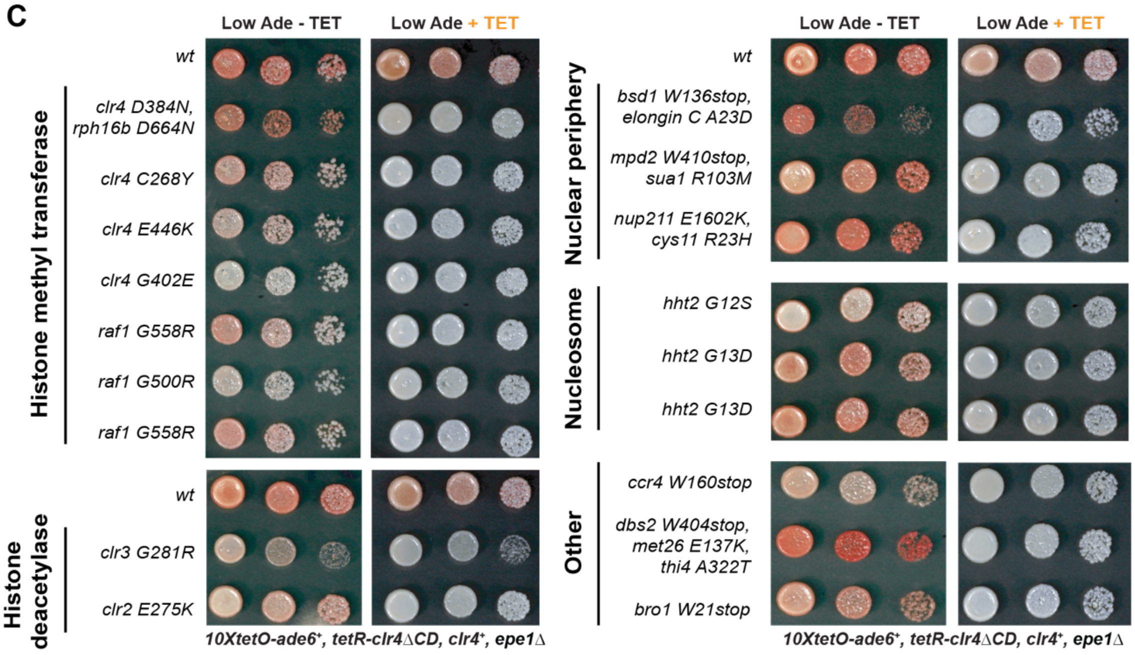

### Figure S2. Shipkovenska et al.

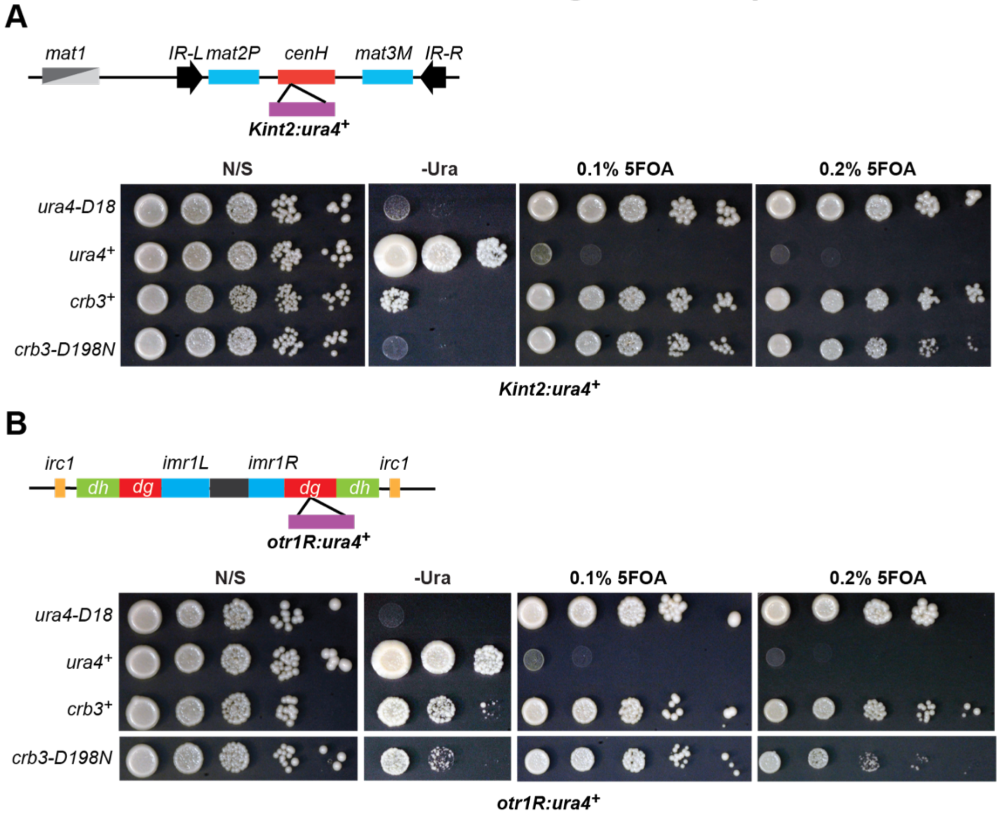

#### Figure S3. Shipkovenska et al.

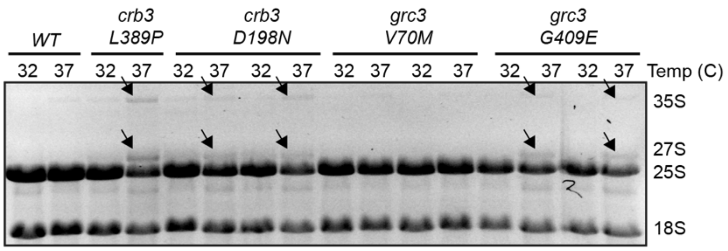

**Figure S4. Shipkovenska et al.**

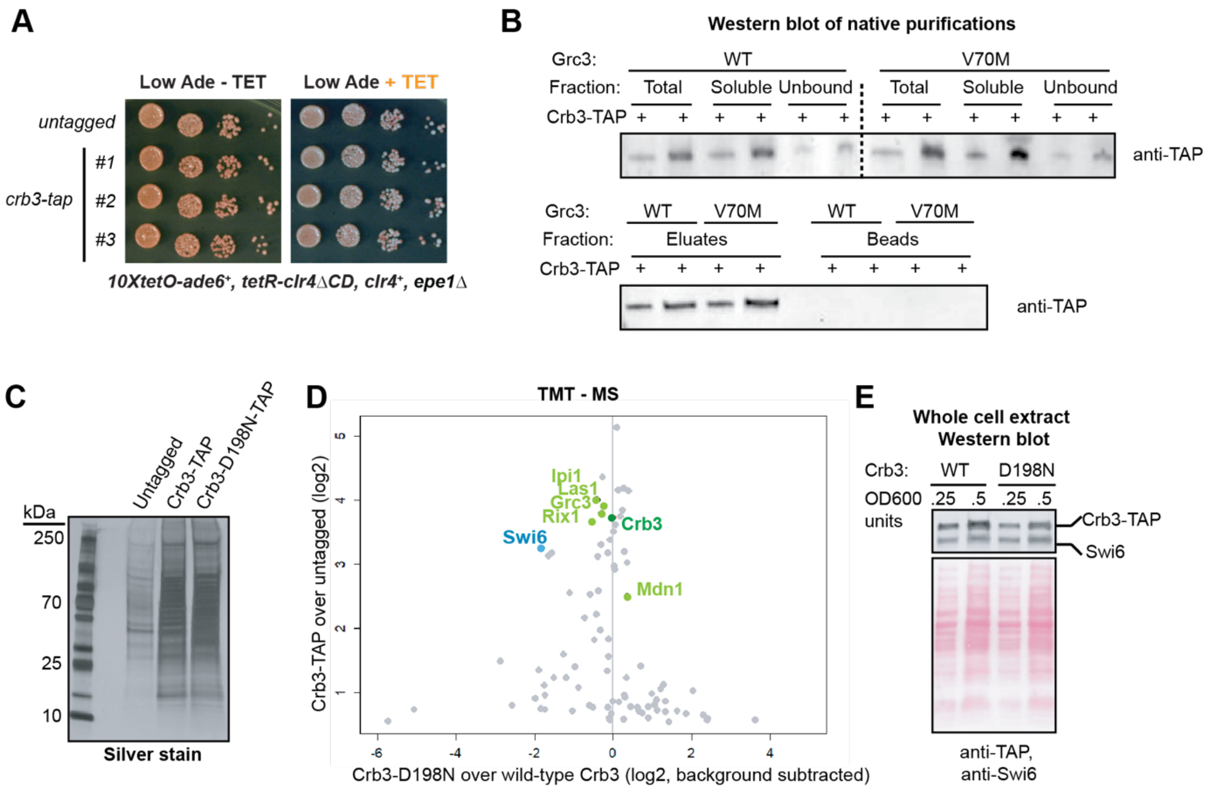

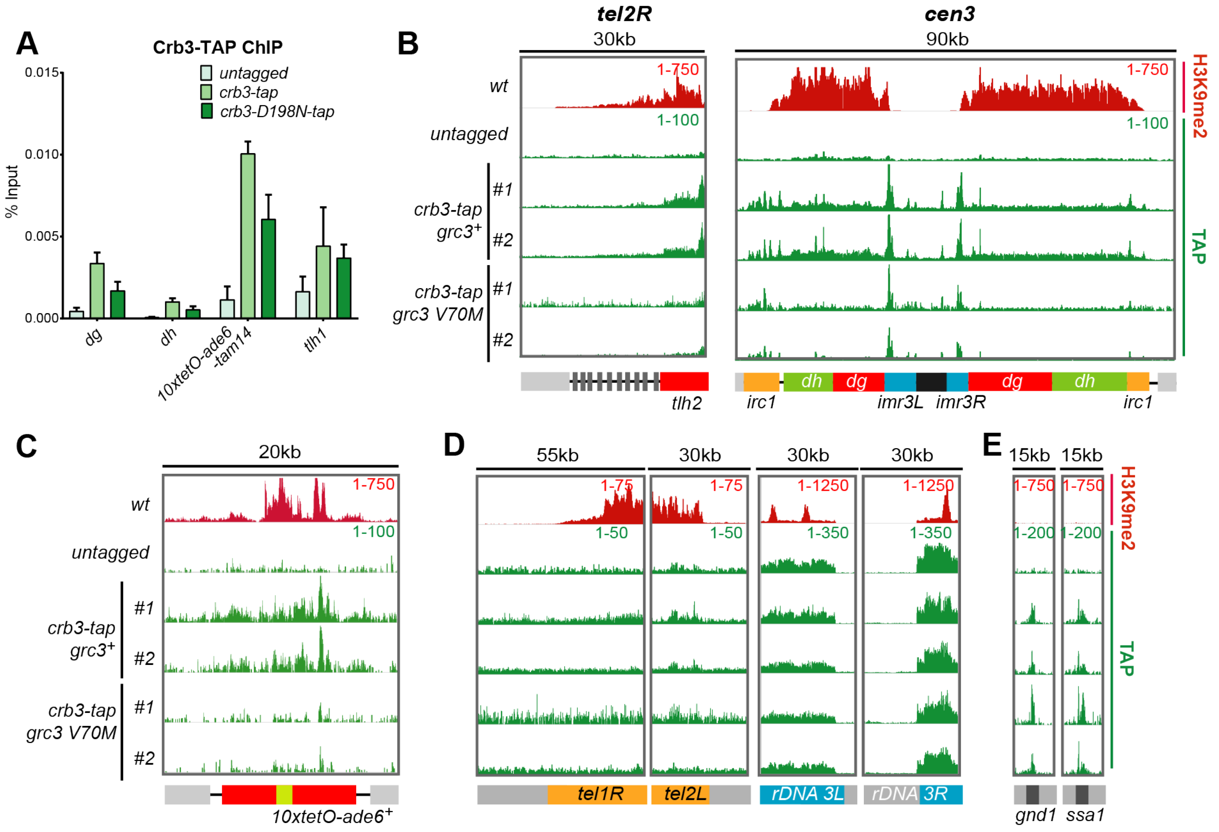
A
Crb3-TAP ChIP
untagged
crb3-tap
crb3-D198N-tap
% Input
0.015
0.010
0.005
0.000
dg
dh
10xtetO-ade6 -tam14
tlh1
B
tel2R
30kb
cen3
90kb
wt
untagged
crb3-tap grc3+
crb3-tap grc3 V70M
#1
#2
1-750
1-100
H3K9me2
TAP
tlh2
irc1
dh
dg
imr3L
imr3R
C
20kb
10xtetO-ade6+
D
55kb
30kb
1-75
1-50
1-1250
1-350
tel1R
tel2L
rDNA 3L
rDNA 3R
E
15kb
1-200
gnd1
ssa1

**Figure S6. Shipkovenska et al.**

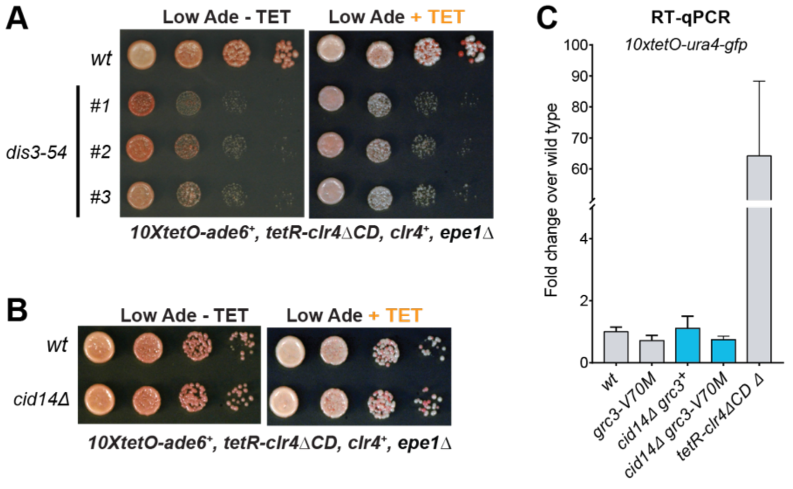

### Figure S7. Shipkovenska et al.

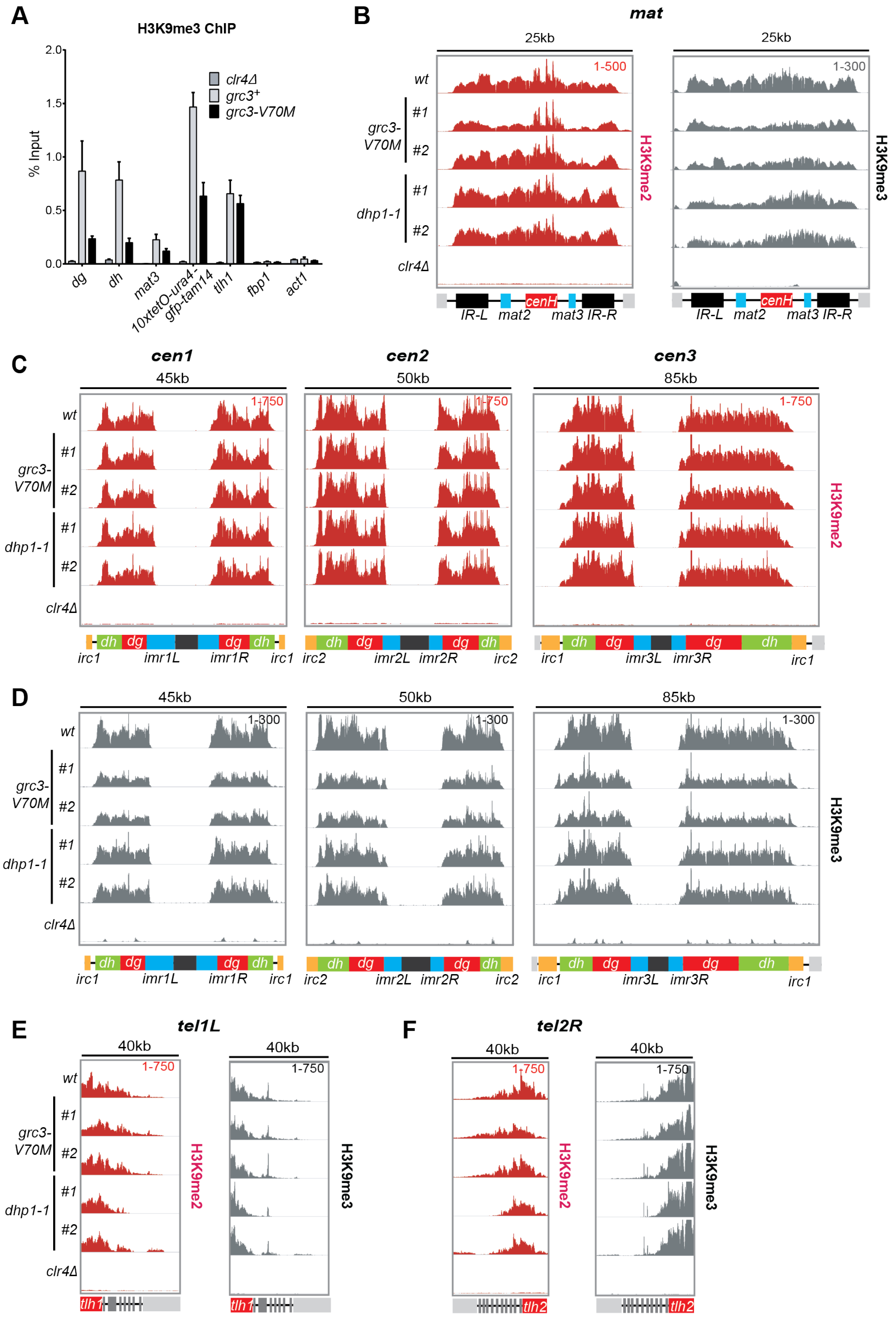

#### Figure S8. Shipkovenska et al.

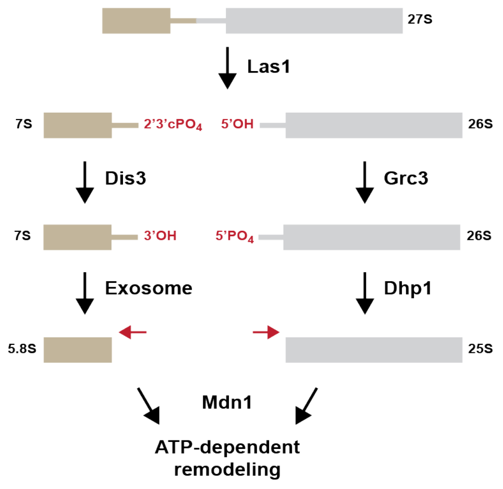
